# On the Spatial Positioning of Ribosomes around chromosome in *E. coli* Cytoplasm

**DOI:** 10.1101/2023.07.04.547709

**Authors:** Abdul Wasim, Palash Bera, Jagannath Mondal

## Abstract

The spatial organization of ribosomes in the cytoplasm of Escherichia coli (*E. coli*) bacteria has been a subject of longstanding intrigue. Previous investigations suggested that ribosomes remain completely excluded from the chromosome-rich nucleoid region in a “poor solvent” like cytoplasm. Here, we present an integrated model of the bacterial cytoplasm, informed by experimental data, which updates this prevailing narrative. We demonstrate that ribosomes maintain a delicate balance of both attractive and repulsive interactions with the chromosome, contrary to the conventional notion of it acting as an inert crowder in cytoplasm. The multi-dimensional spatial distribution of free ribosomes (30S and 50S) and bound ribosomes (70S polysome) inside the cytoplasm reveals that the extent of mutual ribosome-chromosome segregation is relatively less pronounced due to the presence of non-negligible amount of ribosomes within the inner core of the cytoplasm. In particular, we identify a central void within the inner-most core of the nucleoid that lacks chromosomal DNA but can accommodate finite proportion of both free (11 %) and bound (18 %) ribosomes. Furthermore, our analysis of the chromosome mesh size and the conformation of bound ribosomes suggests that bound ribosomes remain elongated and would be able to navigate past the chromosome mesh to access the central void. Together by highlighting the dynamic nature of ribosome localization in E. coli, this investigation proposes that this segregation is crucial for maximizing the utilization of synthesized mRNA and facilitating efficient translation into proteins, which are essential for bacterial survival.

## INTRODUCTION

The archetypal bacteria *E. coli* has a relatively small cell size (approximately 2 *−* 3µm) and possesses a compact genome consisting of 4.64 Mega base pairs (Mbp). Despite lacking the complex compartmentalization found in eukaryotes, the cytoplasm of *E. coli* remains highly intricate due to its in vivo chromosome organization and the interactions between cytoplasmic species and the chromosome. Over the past two decades, significant progress has been made in unraveling the in vivo chromosome organization of *E. coli* through molecular biology techniques[1–3], microscopy studies[4–6], chromosome conformation capture methods[7–9], ChIP-Seq analyses[10, 11], and RNA-Seq investigations[12]. These rapidly advancing datasets, coupled with computational models, have facilitated the elucidation of the architectural characteristics of the chromosome at increasingly higher resolutions, leading to a better understanding of its spatial organization[13–18]. However, a majority of these studies delve into the architecture of the chromosome and the compaction mechanisms driven by various nucleoid binding proteins.

Besides the hierarchical organization of the chromosome, the spatial arrangement of other cytoplasmic particles relative to the chromosome has also been a subject of ongoing investigation[19–21] as it has a potential impact on the bacteria’s survival[22, 23]. Among these particles, the spatial distribution of ribosomes in the cytoplasm and their potential segregation from the chromosome have garnered significant attention[24–27]. Ribosomes, one of the most abundant components in *E. coli* cytoplasm, consist of three subunits: 30S, 50S, and 70S. The 30S and 50S subunits exist as free monomers, while during translation, they combine on mRNA to form the 70S subunit. Each mRNA strand can accommodate up to 13 copies of the 70S subunit, forming a polysome or bound ribosomes. The total number of ribosomes within a cell can exceed 60,000, depending on the growth conditions[28]. In this article, we quantitatively investigate the previous hypothesis regarding the spatial positioning of ribosomes within the *E. coli* cytoplasm using a high-resolution integrative modeling approach.

Experimental investigations using fluorescence microscopy have revealed that ribosomes predominantly occupy the end-caps or poles of bacterial cells, while the chromosome remains at the center[29–31]. Similar observations have been confirmed by electron microscopy experiments on bacteria[32–34]. The segregation of the chromosome and ribosomes in bacteria is thus an important aspect of their survival mechanism. Further exploration using super-resolution microscopy and particle tracking techniques, with *E. coli* as a model organism, has suggested that ribosomes are predominantly excluded from the nucleoid and get localized to the periphery of the cytoplasm[24, 25], as depicted in Figure 1. This exclusion is likely due to repulsion by the densely packed nucleoid, with the expelled ribosomes acting as macromolecular crowders for the chromosome[19, 35, 36], creating an environment in which the cytoplasm is a poor solvent for the chromosome[37]. Previous studies on the spatial positioning of ribosomes have traditionally treated them as inert crowders, assuming that ribosome-DNA interactions are solely driven by non-specific excluded volume effects[14, 38–41]. This has led to the long-held notion that the chromosome and ribosomes are spatially anticorrelated and completely segregated from each other. However, recent findings from linear density profiles of the chromosome and ribosomes across the cytoplasm[27] have demonstrated substantial densities of bound ribosomes (polysomes) as well as free ribosomes within the nucleoid-containing region of bacterial cells.

**FIG. 1.**
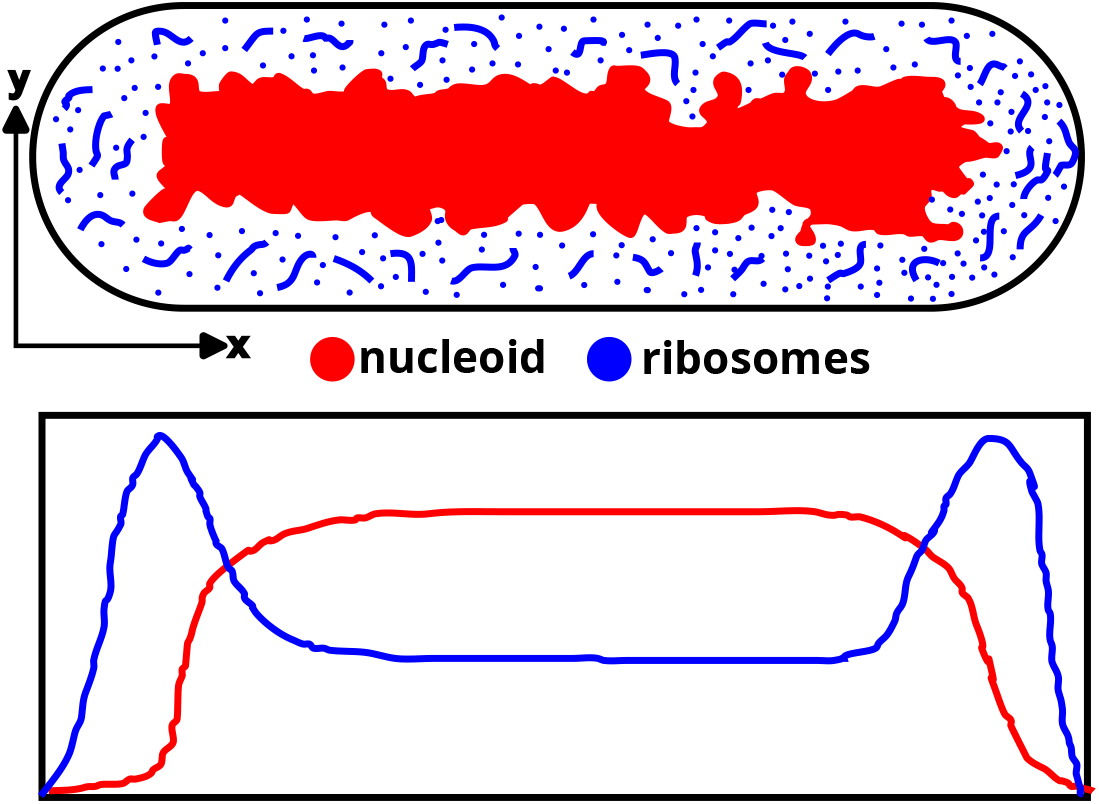
The schematic depicts the spatial segregation of the ribosomes and the nucleoid as described by experiments. A representative linear density has been shown below the picture of the cell.

In this study, we utilize a recently developed integrated model of the *E. coli* cytoplasm[18], which incorporates a data-rich model of the chromosome at a resolution of 500 base pairs (bp) and optimized the chromosome-ribosome interactions to reproduce experimentally reported chromosome and ribosome axial densities[27]. We aim to shed light on the specific interactions between ribosomes and the chromosome, challenging the notion that ribosomes are inert crowders. Through the analysis of two and three dimensionally resolved densities of the chromosome, free ribosomes, and bound ribosomes, we provide a comprehensive characterization of the chromosome-ribosome segregation[24, 25, 27]. Notably, our investigation reveals the presence of a void in the central nucleoid region and identifies a significant population of both free and bound ribosomes in this central location, in addition to their usual location at the periphery.

## RESULTS

In the current work, we explore the distribution of monomeric ribosomes (referred here as ‘free ribosomes’) and polysomes (referred here as ‘bound ribosomes’) inside *E. coli*. For this purpose, we simulate a minimal model of cytoplasmic environment of the bacteria in a data-informed fashion. We utilized a spherocylindrical confinement that matched the dimensions of bacterial cells in a medium growth condition (specifically, wild-type *E. coli* cells grown in M9 minimal media at 30C). Within this confinement, we developed a high-resolution hyper-branched model of the *E. coli* chromosome. This model incorporated various experimental data sources, including Hi-C and RNA-seq data, as well as multiple other experimental datasets[1, 12, 42–45]. To generate chromosome conformations, we employed a polymer framework that represented each bead as 500 base pairs. The details of the chromosome model can be found in our recent publication[18], and a brief description is provided in the *Methods* section. Around the chromosome, we pack the cytoplasm with bound and free ribosomes with the number, their sizes and masses being commensurate with the growth condition (Table-I)[14, 46, 47]. Here we investigate the distribution of the ribosomes inside the cell and we try to address the discrepancies and reconcile both conflicting ideas of ribosomes exclusion from nucleoid and ribosome occupancy inside the nucleoid through our data-informed model, which has been made available at the following GitHub repository: https://github.com/JMLab-tifrh/ecoli_finer.

**TABLE I.** Summary of the incorporation of free and bound ribosomes

| subunit | number of subunits | size (nm) | mass (MDa) |
| --- | --- | --- | --- |
| 30S | 1292 | 14 | 0.84 |
| 50S | 1292 | 17 | 1.36 |
| 70S | 18416 | 21 | 2.3 |

\*a free:bound = 20:80 has been considered for this study since we
have simulated a WT *E. coli* cell grown in M9 minimal media at 30C

### Ribosomes are not just *inert* crowders for the chromosome

Ribosomes are one of the most abundant moieties in the cytoplasm of *E. coli*, consisting of free monomeric species (30S and 50S) and bound ribosomes, which are 13-meric chains of 70S ribosomal subunit. As illustrated in Figure 2a, in present work monomeric 30S and 50S have been modeled as ‘free’ spherical particles with masses and sizes equivalent to experimental report[24, 47] (Table-S1). On the other hand, the bound ribosomes have been modeled as polymeric chains of thirteen copies of 70S ribosomal subunit, ‘bound’ to each other. In our simulation of an *E. coli* cell under moderate growth conditions (M9 minimal media at 30C), we incorporated a total of 21,000 ribosomes[46]. Based on our consideration of these growth conditions, we assume that 80% of the ribosomes are actively translating, meaning they are “bound” to mRNA and involved in protein synthesis. The remaining 20% of ribosomes are non-translating, exist as monomers, and are evenly distributed between the 30S and 50S subunits[14, 27, 48].

**FIG. 2.**
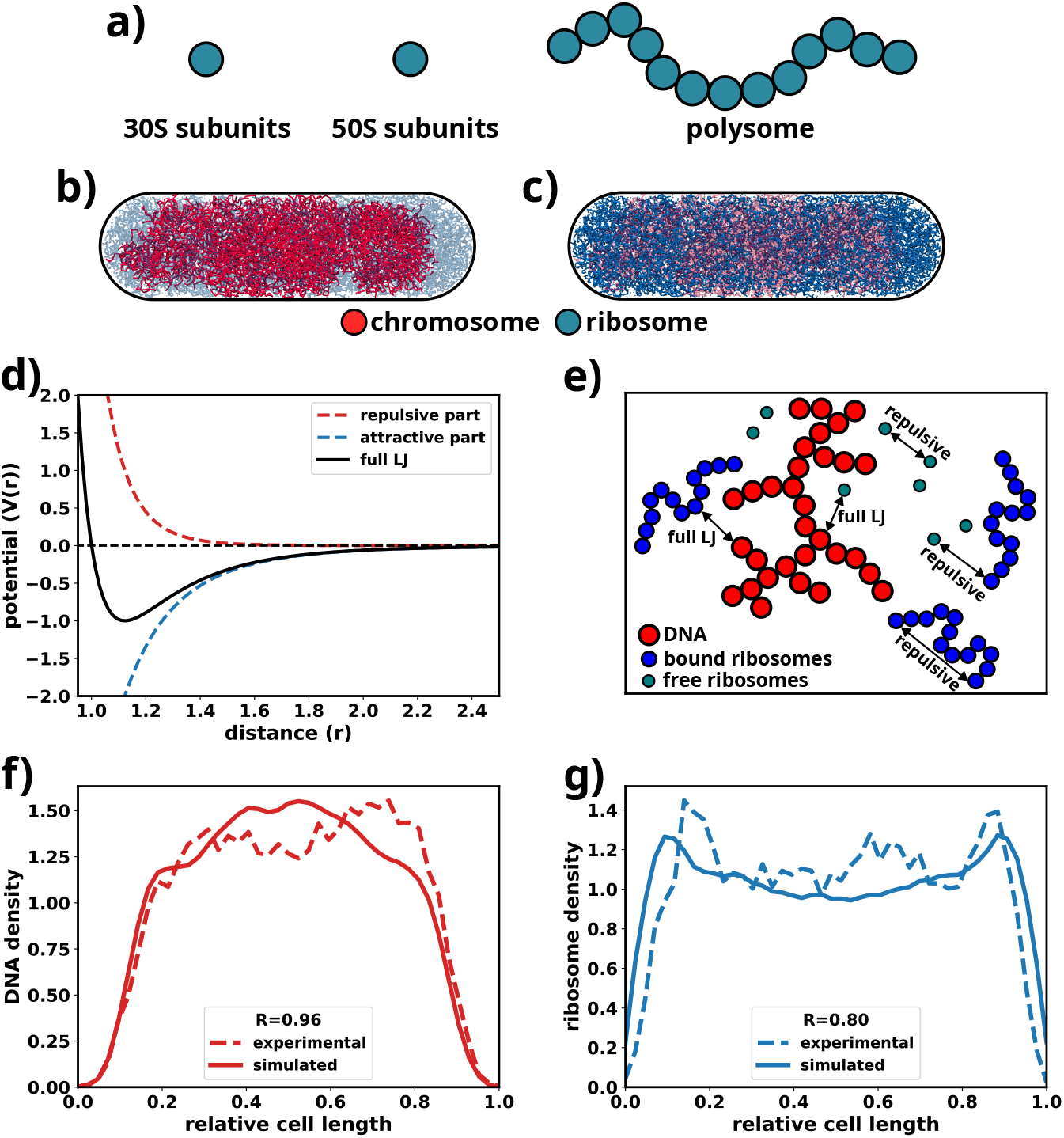
**a)** A schematic showcasing how different ribosomes have been modeled. 30S and 50S have been modeled as spherical particles while bound ribosomes have been modeled as a 13-mer of 70S subunits. Each 70S subunit has been considered as a spherical particle. **b)** A representative snapshot showing the chromosome in red. The ribosomes have been faded out to highlight the chromosome. **c)** A representative snapshot showing the ribosomes in blue. The chromosome has been faded out to highlight the ribosomes. **d)** A depiction of the repulsive (red), attractive (blue) parts of the Lennard-Jones potential (black). **e)** A schematic of the various non-bonded interactions present in the model. **f)** A comparison of simulated and experimental chromosome linear densities for the optimized model. **g)** A comparison of simulated and experimental ribosomes linear densities for the optimized model.

In majority of the precedent modeling attempts ribosomes and chromosome for an *E. coli* cell[14, 24, 40, 41], ribosomes have been traditionally treated as *inert* crowders, devoid of any specific attractive interaction with the chromosome. Majorly driven by qualitative hypothesis[24, 25] that nucleoid and ribosomes mutually exclude each other in *E. coli* cytoplasm, DNA-ribosome interactions had previously been modeled via purely excluded volume interaction (which are essentially soft repulsive interaction between DNA and ribosomes). In the present investigation we depart from this notional view of excluded volume interaction and opt for a data-informed approach by deriving chromosome-ribosome interactions by directly benchmarking against experimentally measured axial density profiles of chromosome and ribosome[27]. When compared with experimentally reported density profile, we find that a purely excluded volume interaction between ribosome and chromosome would over-amplify the anti-correlated nature of ribosome and chromosome densities (Figure S1). Towards this end, we employ a functional form that consists of both the attractive and repulsive parts, to model the chromosome-ribosome interactions and then iteratively optimize the interaction parameters via series of simulation (see *Methods* and Figure S2 and Figure S3) until a good fit between the simulated and experimental linear densities are achieved. The interaction potentials for each ribosome subunit and the chromosome has been shown in Figure S4.

Figure 2b shows the DNA in red and the ribosomes (30S, 50S and 70S) in light blue while Figure 2c depicts the ribosomes in blue and chromosome in light red. The functional forms of the various interactions such as the DNA-ribosome interactions have been showcased in Figure 2d. A schematic showing how different components of the model interact has been shown in Figure 2e which highlights the fact that the interactions of DNA with the ribosomes have been modeled using both attractive and repulsive parts. The interaction parameters, namely A (determines the strength of the repulsive part) and B (determines the strength of the attractive part) have been optimized by considering the Pearson correlation coefficient (R) between linear densities calculated from the simulations with that of the experiment, as can be seen from Table-II.

**TABLE II.** Variation of R with A and B for DNA and bound 70S subunit interaction

| **A** | **B** | **Comment** | **$R_{DNA}$** | **$R_{rib}$** |
| --- | --- | --- | --- | --- |
| 1.00 | 0.00 | purely repulsive | 0.68 | 0.08 |
| 0.50 | 0.10 | weakly attractive | 0.88 | 0.82 |
| 0.25 | 0.01 | moderately attractive | 0.98 | 0.83 |
| 1.00 | 1.00 | strongly attractive | 0.78 | 0.15 |

\*\*A and B are the coefficient of the repulsive and attractive parts of the Lennard-Jones function in Figure
2d. $R_{DNA}$ and $R_{rib}$ are the Pearson correlation coefficients between experimentally measured and simulated linear densities for DNA and ribosomes, respectively. Optimum values of A and B have been provided in Table-S1.

After successful optimization, the corresponding simulated linear density of the chromosome has been depicted in Figure 2f by the bold red line. The respective experimental linear density has been represented as the dashed red line. The experimental and the simulated densities share a Pearson correlation coefficient (R*_DNA_*) of 0.96 which shows that the simulations using the current interaction parameters can recapture the experimental data to a high degree of accuracy.

From Figure 2c, it can be seen that the ribosomes tend to flock at the end caps of the cell. It can be verified by the linear density plots shown in Figure 2g where the simulated ribosome linear density has been represented as the bold blue line while the experimental linear density has been represented by the dashed line. The experimental and simulated linear densities have a Pearson correlation coefficient (R*_rib_*) of 0.80 which signifies a high degree of reproducibility of the experimental data. The peaks on both ends of the linear density showcase the higher propensity of ribosomes to populate the ends of the cell. The analysis together suggests that ribosome is just not an inert, non-interacting crowder, as had been traditionally posited in previous investigations[14, 40, 41]; rather a fine balance between attractive and repulsive interaction between ribosome and chromosome provide a quantitatively more accurate description of chromosome-ribosome spatial positioning. As would be elaborated in the upcoming sections, the requirement of presence of both attractive and repulsive interaction between DNA and ribosomes is also suggestive of non-negligible presence of ribosomes inside chromosome mesh, in contrast to long-held perspective of complete exclusion of ribosome by DNA.

### The model cytoplasm represents a poor solvent with a *∼*50 nm mesh size

The relative compaction of a polymer in a solvent is an interplay of the intra-polymer and polymer-solvent interactions. This interaction is characterized by the Flory exponent ν. The Flory exponent is a parameter that defines how good the solvent is for a polymer (here chromosome). If ν < 0.5, then the solvent is a poor solvent and hence the polymer would form compact, collapsed conformations. For ν = 0.5, the polymer behaves like a randomwalk polymer or a polymer in θ solvent. For ν > 0.5, it swells up and forms loose, extended conformations. The scaling exponent for the *E. coli* chromosome was recently reported to be *∼*0.36 which indicates that inside the cell it behaves like a polymer in a bad solvent[37]. As we have modeled the cytoplasm explicitly while using the experimental linear densities as a benchmark, we wanted to calculate the Flory exponent from our simulations itself.

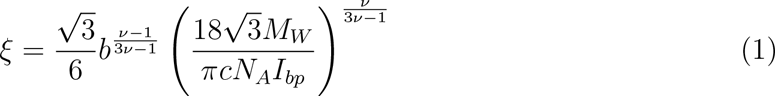

Eq-1 was used by Xiang et al.[37] to calculate the value of the mesh size (ξ) from experiments. We used the same equation to estimate the value of ν from it (Eq-1). To this end we calculated the mesh size (ξ), the Kuhn length of the chromosome from our polymer model (b), the molecular weight of 500 base pairs of chromosome (M*_W_*), which is equivalent to a single polymer bead in the model, concentration of DNA within the volume occupied by the chromosome (c) and the size of a base-pair of DNA (I*_bp_*). The values we obtained from the simulations or literature for the parameter set are as follows: ξ *∼* 50 nm, c = 9.5 mg/ml, b=63.32 nm, M*_W_* =650 g/mol and I*_bp_*=1 nm[49, 50] (see *Supplementary Results*). Incorporating all these values into Eq-1 yields the result ν *∼* 0.31 (Figure S5) which corroborates well with the experimentally reported value. This showcases that the chromosome-ribosome interactions used in our model are able to capture the poor nature of the cytoplasm as solvent for the chromosome. Moreover, a Flory exponent of ν *∼* 0.31, which matches well with the experimentally determined value of ν *∼* 0.36[37], signifies that the cytoplasm acts as a poor solvent for the chromosome. This results in the chromosome being able to exist in a “compact” conformation (Figure 2b), commensurate with poor solvent quality of cytoplasm.

### The chromosome conformation is not significantly affected by the free to bound ribosome ratio

We next wanted to investigate how the presence of different amounts of monomeric ribosome (30S/50S) (i.e. ‘free ribosome’) and polysomal ribosomes (70S) (i.e. ‘bound ribosome’) affect the chromosome conformation. To that end, we simulated a diverse realization of cytoplasmic composition where we varied the bound to free ribosome ratios (retaining same chromosome model). In particular, we individually simulated bacterial cytoplasm with following 70S:30S:50S ratios: 0:50:50, 20:40:40, 50:25:25, 80:10:10 (original and majorly used in this study) and 100:0:0. We calculated the linear density profiles for the chromosome and ribosomes for all these compositions (Figure 3).

**FIG. 3.**
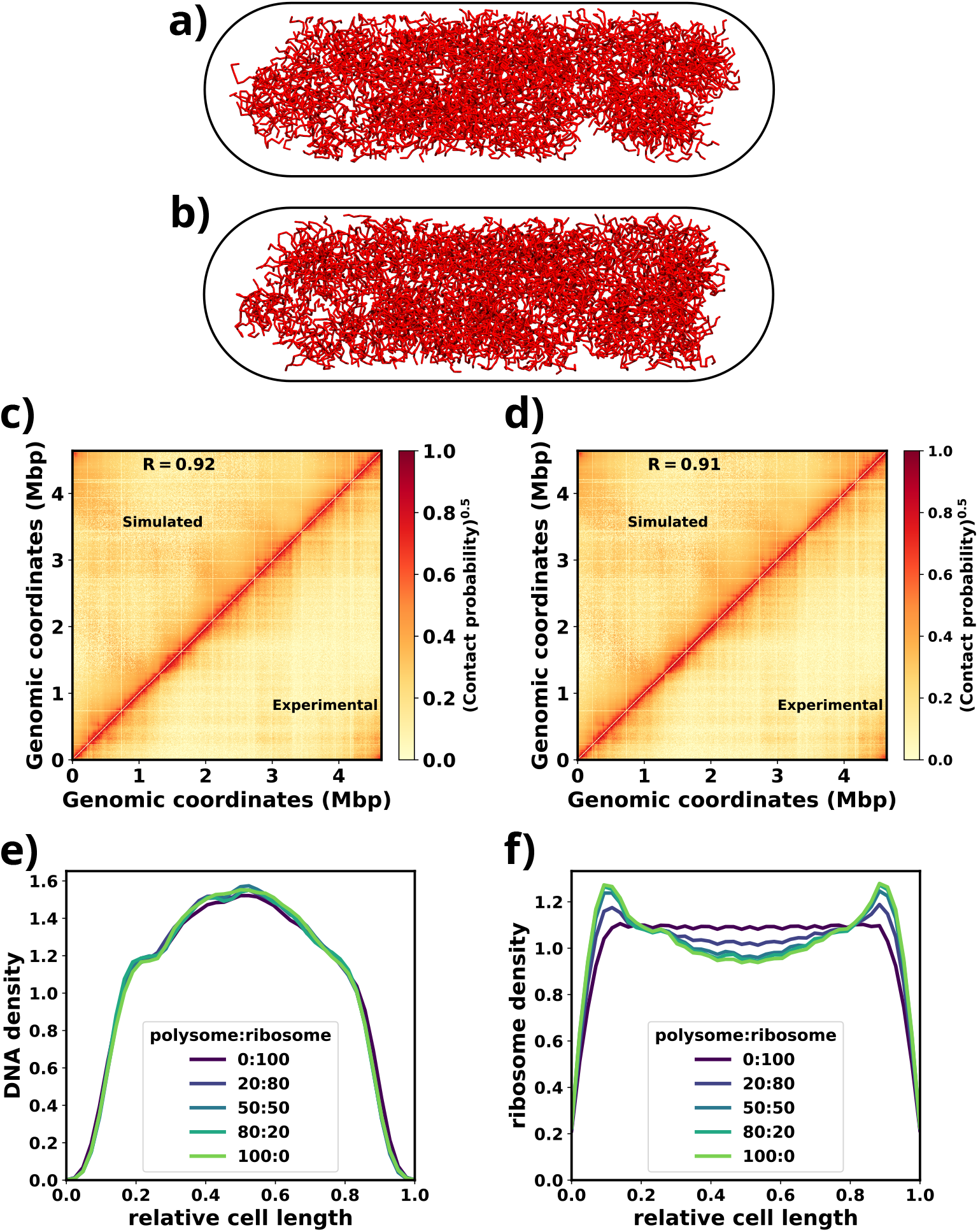
**a)** Chromosome conformation for 0:100 bound:free ribosome ratio. **b)** Chromosome conformation for 100:0 bound:free ribosome ratio. **c)** Comparison of intra-genome contact probability matrix with the experimental Hi-C contact probability matrix for 0:100 bound:free ratio. **d)** Comparison of intra-genome contact probability matrix with the experimental Hi-C contact probability matrix for 100:0 bound:free ratio. **e)** Linear density of chromosome for varying bound to free ribosome ratios. **f)** Linear density of ribosomes for varying bound to free ribosome ratios

Figure 3a and Figure 3b demonstrate chromosome conformations for two extreme scenarios: a) where all are free ribosomes, i.e. no bound ribosomes are present. b) all are bound ribosomes, i.e. only bound ribosomes. Both these conformations appear to be very similar which has been reflected in Figure 3c and Figure 3d. Both Figures 3c and d show that for both scenarios a) and b), the simulated intra-genome contact probability map does not change significantly. Thus we think that the chromosome conformation is unaffected by the amount of bound or free ribosomes present.

We can also see from the Figure 3e that the chromosome linear densities do not change significantly with respect to variation of bound:free ribosome ratios. However from Figure 2f, it can be seen that when the cytoplasm is filled only with free ribosome (bound:free = 0:100), the linear density of the ribosome is almost flat in the nucleoid region (0.2 to 0.8 of the relative cell length). As the fraction of bound ribosomes gradually increases, the linear density of ribosome on the nucleoid region slowly decreases to a minimum in a cytoplasm containing only ‘bound’ ribosome (bound:free ribosome ratio 100:0). This suggests that bound and free ribosomes are differentially excluded from the region where the nucleoid resides: free ribosomes can, to a reasonable extent, populate the nucleoid region while the bound ones are predominantly excluded out form the nucleoid. However it should be noted that for none of bound:free ribosome ratios, does the linear density fall to very low values in the nucleoid region. Therefore, even in the case of the cytoplasm filled with 100% bound ribosomes, linear densities indicate finite presence of bound ribosomes inside the nucleoid populated region. Though the consensus hypothesis is that bound ribosomes are excluded out from the nucleoid, a similar observation, that bound ribosome densities do not fall to really low values, can be made from even the experimental linear densities[19, 25]. Realizing that the linear densities alone are not sufficient to fully resolve the exclusion of bound ribosomes from the nucleoid, we decided to investigate the spatial positioning of free and bound ribosome via characterizing their spatial density maps.

### Spatial density maps reveal the presence of a ribosome populated central void

To resolve the extent of spatial segregation of bound and free ribosomes from the nucleoid and especially to reconcile the observed presence of free ribosome inside nucleoid-rich location, we calculated the spatial (number) density map of DNA and all ribosomes (both “free” and “bound”) along the cross section of the bacterial cell (as shown in Figure 4a). A representative conformation of the DNA viewed from the cross-section axes has also been shown in Figure 4b which hints at a possible low density region at the center. The two-dimensional DNA density (σ*_DNA_*) shown in Figure 4c aptly captures such an observation. We can observe from Figure 4b that the inner core of the central regions of the cytoplasm has very low DNA density. Encircling it is a region of high DNA density, beyond which the DNA density wanes out till the periphery. The low DNA density at the centre indicates the presence of a void at the centre along the long axis of the cell.

**FIG. 4.**
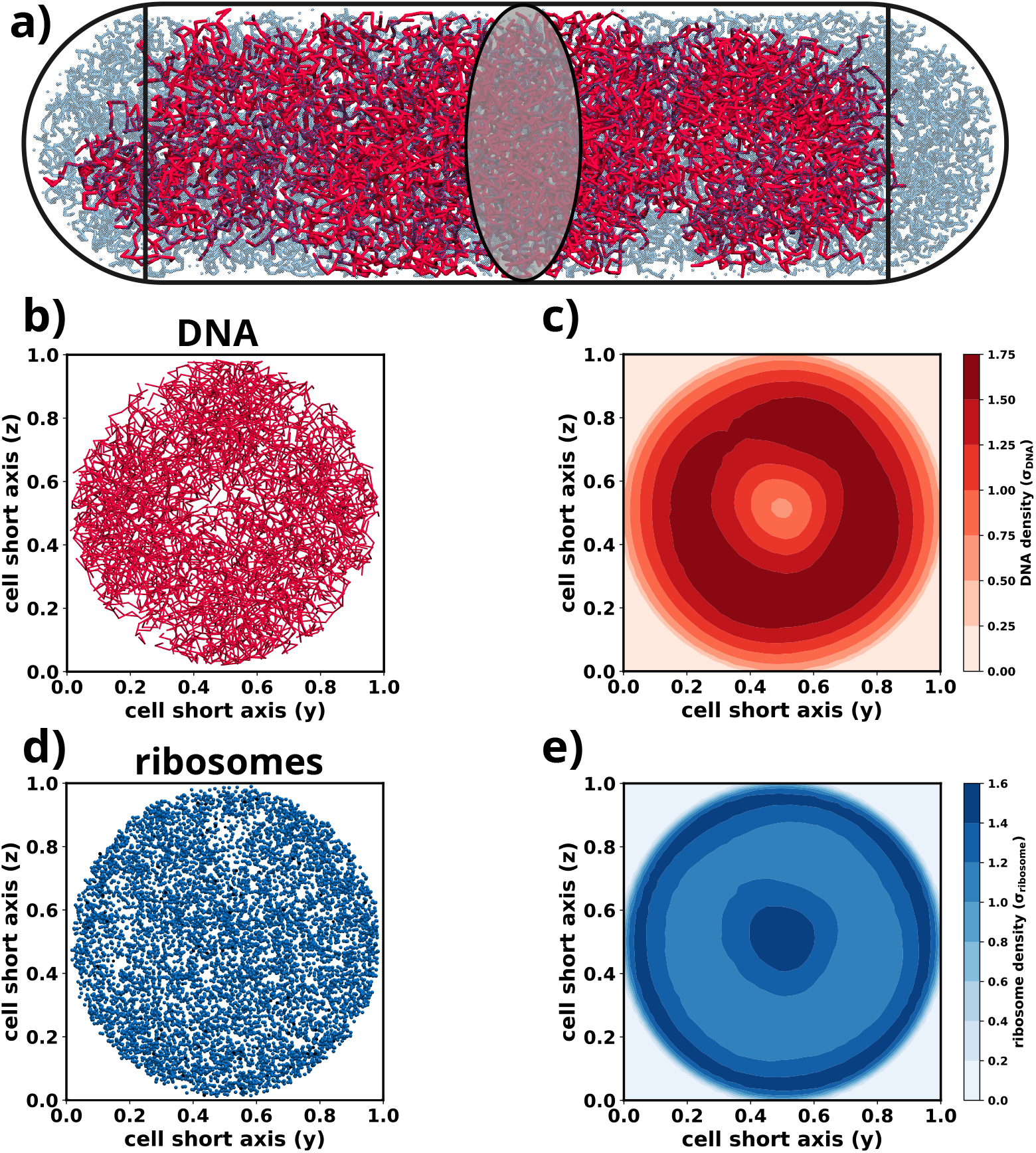
**a)** A snapshot of the chromosome showing the cylindrical region and a cross-section of the cell along which the average 2D densities have been calculated. **b)** A snapshot of the DNA along the cross-sectional axes. **c)** Contour plot showing the DNA density (*σ_DNA_*) along the radial directions. **d)** A snapshot of the ribosomes along the cross-sectional axes. **e)** Contour plot showing the ribosome density (*σ_ribosome_*) along the radial directions.

Although it is difficult to understand the distribution of the ribosomes along the cross-sectional axes from the representative conformation in Figure 4d, the ribosome density map(σ*_ribosome_* in Figure 4e) shows complementarily opposite localization trend as that of DNA. At the centre, where there is low DNA density, there is a very high ribosome density. The next concentric region, which is highly populated by DNA, has very low ribosome density followed by high ribosome density near the periphery. Thus ribosome and DNA localization are anti-correlated. However, as a key observation, we do see presence of ribosomes at the centre, in the cavity, which is buried inside the nucleoid[25, 27].

To confirm that the central void is not an artifact of computational architecture, we performed multiple control simulations such as i) simulating the chromosome in absence of ribosomes to prove that the void is an intrinsic feature of the chromosome. ii) generating the initial configuration for the simulations via an alternate method to show that it is not a detect of initial conformation generation. iii) using purely inert crowders (via employing repulsive DNA-ribosome interaction) to show that the appearance of void does not depend upon the chromosome-ribosome interaction (see *Supplementary results*). These control simulations validates the presence that the void inside the nucleoid.

Interestingly the structures obtained from a different model by Hacker et al.[15] revealed the presence of voids inside the nucleoid. Thus the central pore we observe from our model conforms to the previous observations regarding the pore(s) or voids inside the nucleoid and further provides confidence regarding the observation of the central void in simulations.

### The central void is DNA free and is populated by free and bound ribosomes

To quantitatively observe the presence of ribosomes inside the void at the center of the nucleoid, we first render a contour plot of the spatial difference in the densities of DNA and ribosomes, as shown in Figure 5a. The blue regions of the heatmap have more ribosomes density than the chromosome while regions which are red have more DNA density as compared to the ribosomes. White regions indicate similar DNA and chromosome densities. Therefore the heatmap, having a blue region at the centre, points to the fact the void inside the nucleoid is populated by ribosomes.

**FIG. 5.**
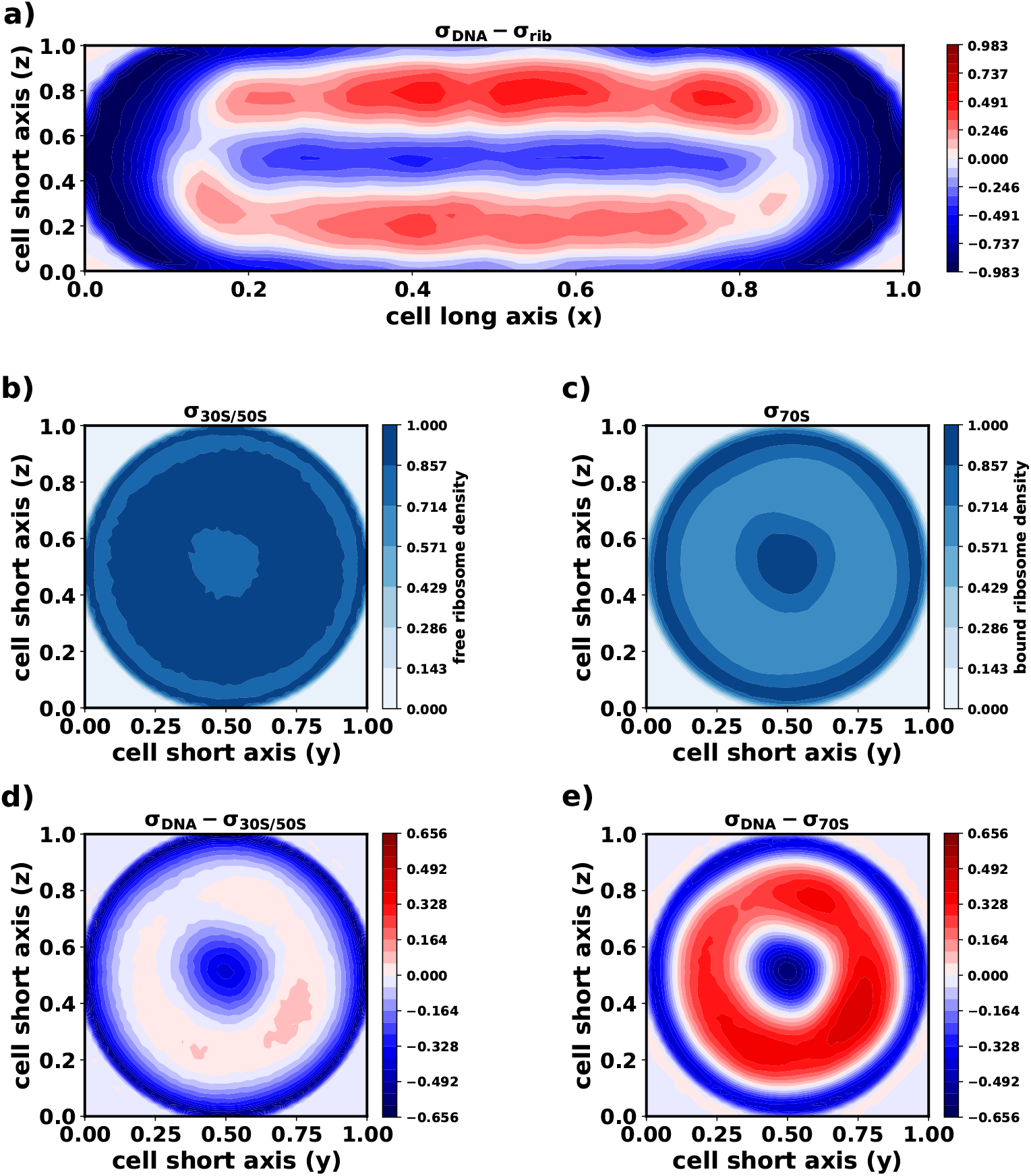
**a)** Contour plot showcasing the differences in the densities of the DNA and the ribosomes in the cross-section of the cell with the axial and one short axis (xz-plane). The central region in blue showcases the void. **b)** Contour plot showing the free ribosome density (*σ*_30_*_S/_*_50_*_S_*) along the radial directions. **c)** Contour plot showing the bound ribosomes density (*σ_polysome_*) along the radial directions. **d** Contour plot showing the difference in free ribosome and DNA density (*σ_DNA_*-*σ*_30_*_S/_*_50_*_S_*) along the radial directions. **e)** Contour plot showing the difference in bound ribosomes and DNA density (*σ_DNA_*-*σ*_70_*_S_*) along the radial directions.

To understand if the distribution of ribosome is contingent on their inherent nature (‘free’ versus ‘bound’), we plot the spatial heat maps for free (30S/50S) (Figure 5b) and bound (70S) (Figure 5c) ribosomes, individually. Very interestingly, we see that the central region is occupied by both bound and free ribosomes. For a comparative visualization of DNA vs free/bound ribosome densities, we plot the differences in densities between DNA and monomeric ribosomes in Figure 5d and between DNA and bound ribosomes in Figure 5e respectively. In these figures, red means the region has a higher DNA density than that of the ribosome. Blue means the region has a higher ribosome density. White signifies similar DNA and ribosome densities. We see that in Figure 5d that free monomeric ribosomes do occupy regions not occupied by the DNA (blue regions at the centre and the periphery). But it can also co-exist in the chromosome mesh as conveyed by the nearly white region in between the blue regions. On the other hand, from Figure 5e, it can be seen that DNA excludes polysome or bound ribosomes out from the regions it occupies as depicted by concentric blue and red bands. Thus we conclude that DNA excludes out bound ribosomes as previously seen via experiments[19, 27] and previous theoretical studies[14]. This exclusion of the bound ribosomes can be towards the cell periphery or could be also towards the void that has been shown to exist at the centre of the nucleoid. Therefore for the later sections, we consider the void to be also ‘outside’ the nucleoid.

One reason for the free ribosomes not being excluded out of the nucleoid is their relatively small size, with respect to the bound ones. This allows them to freely pass through the chromosome mesh of *∼*50 nm and not being excluded out. Therefore the presence of the free ribosomes in the central void is not surprising and to some extent trivial. However, the presence of bound ribosomes inside the void is interesting. Axial linear densities, which have been used previously to explore the chromosome-ribosome segregation, showed a significant amount of ribosome density in the nucleoid region[19, 25, 27]. The presence of a void, such as this, reconciles with the appearance of the high linear density of ribosomes in the region where the nucleoid resides. 2D heatmaps provide conclusive evidence that free ribosomes are not excluded from the nucleoid, while bound ribosomes are excluded from the nucleoid. However, this gives rise to an important question: if bound ribosomes occupy the “buried” void but are excluded from the nucleoid, can bound ribosomes, near the cell wall, gain entry to the central pore? We try to address this in the next section.

### Bound ribosomes can percolate through the mesh to gain access to the central void

In the previous section we saw that ribosomes occupy the central region of the bacterial cell, encompassed by the DNA in a concentric manner. We have also estimated that the chromosome is organized in such a manner that it creates a mesh with a pore size of about 50 nm. Therefore we wanted to understand, despite having a polymeric architecture, how the bound ribosomes at the surface would be able to penetrate the chromosome mesh and access the central void. Towards this end, we analyze a few representative polysome conformations (see *Supplementary Results*). They show that the bound ribosomes remain mostly in their extended states. Since the bound ribosomes are present in extended conformations, the effective cylindrical volume for them would provide an estimate if they can percolate through the mesh of size 50 nm. Therefore we determined the dimensions of the cylinder that would encompass the conformations (Figure 6a).

**FIG. 6.**
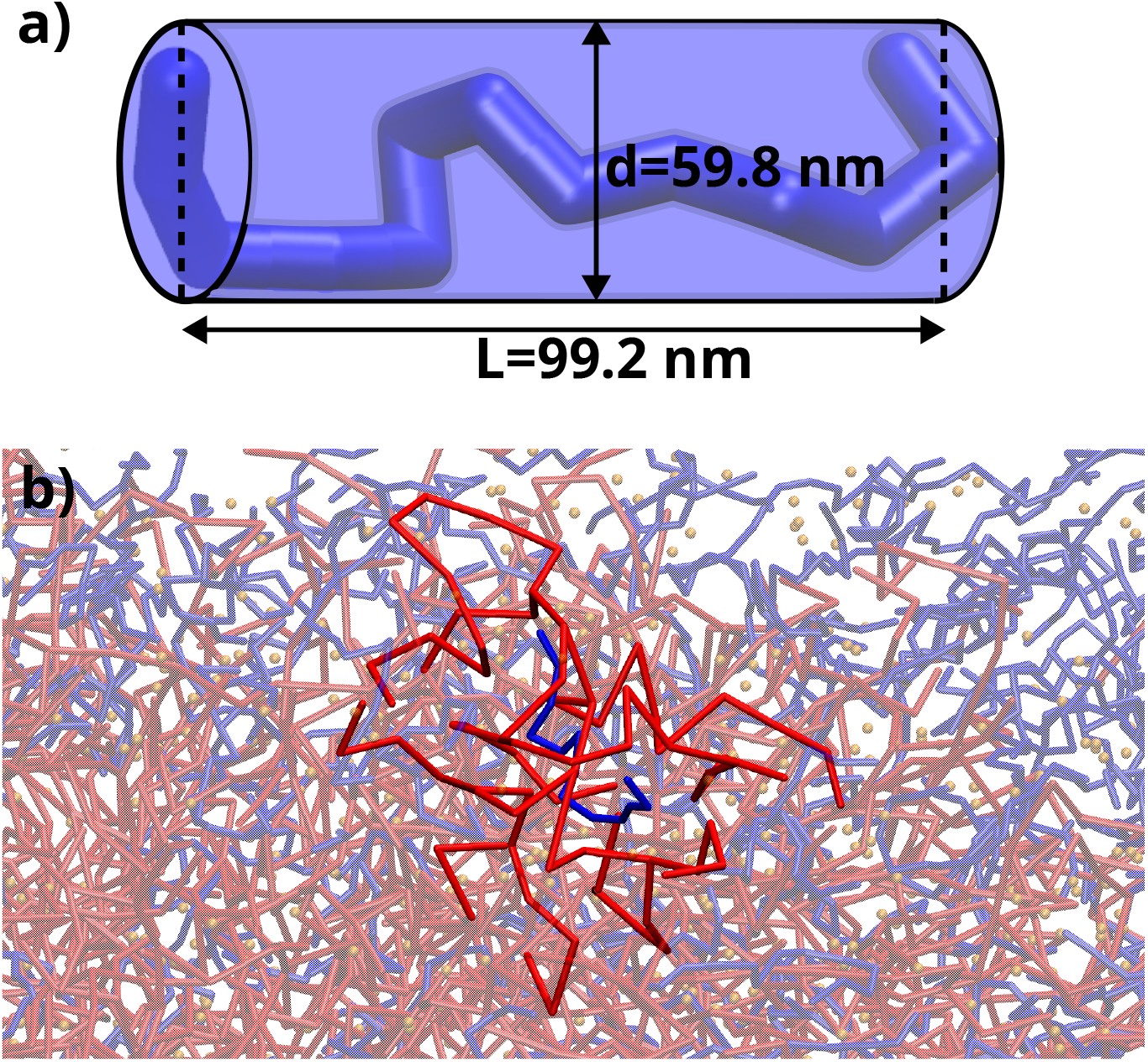
**a)** A polysome conformation estimated via a cylinder. d is the diameter of the cylinder and L is its length. **b)** a representative snapshot showcasing a polysome (blue) passing through the chromosome mesh (red).

We determined that the length of the cylinder is *∼*100 nm while the diameter is *∼*60 nm. Thus the breadth of the cylinder is very close to the size of the mesh created by the chromosome (*∼*50 nm). Taking into account the fact that ribosomes are actually not a rigid cylinder but is a flexible polymer, they would easily percolate through the chromosome mesh and populate the inner void as can be seen from Figure 6b. To conclusively prove that, we ran a set of control simulations (see *Supplementary Results*) where all the bound ribosomes were kept outside the nucleoid, initially (Figure S6), and the simulations were performed to equilibrate the system. As can be seen from Figure S7 that the chromosome and the ribosomes linear densities obtained from the simulations are same as that of Figure 2c and 2e. Thus we conclude that the polysome can percolate through the chromosome mesh, via a worm like movement, to gain access to the central void and reside there. It should be noted that this does not mean that the bound ribosomes are not excluded out of the nucleoid. This only means that the nucleoid accommodate a finite amount of the bound ribosomes due to its mesh like structure and we think that this could be the reason why a very tiny fraction (*∼*8%) of the bound ribosomes were not excluded out of the nucleoid, as can been seen from experimental results[27].

## DISCUSSION

The present study provides compelling evidence that ribosomes play an active role beyond serving as inert cytoplasmic crowders. By comparing experimentally derived axial density profiles of the chromosome and ribosomes, we demonstrate a specific and finely balanced interaction between them. This observation supports previous hypotheses proposing the presence of expanding and contracting forces within the bacterial cytoplasm, which contribute to spatial organization[51].

Through meticulous spatial analysis, we uncover a nucleoid-free void at the center of the cytoplasm that is inhabited by both free and bound ribosomes. To ensure the validity of our findings, we conduct a series of rigorous control simulations, confirming that the observed void is not an artifact of our computational architecture. Notably, we find a substantial density of polysomes within this void, prompting us to investigate whether bound ribosomes can effectively traverse the chromosome mesh and access the void. Our findings reveal that bound ribosomes employ a worm-like movement to navigate through the chromosome mesh, allowing them to gain access to the void.

Furthermore, we quantitatively assess the distribution of ribosomal subunits within the nucleoid-contained region, specifically the cylindrical part of the cell, as well as the portion excluded from it. This analysis is based on comprehensive 3D number densities of ribosomes and the chromosome.

Our observations provide a reconciliation for previous experimental findings[27] indicating that a small fraction of bound ribosomes is not excluded from the nucleoid. There appears to be a partitioning of bound ribosomes, with the majority being excluded from the nucleoid while a subset remains inside. To determine the proportions of free and bound ribosomal subunits that are excluded from the nucleoid, we calculated the percentage of ribosomes residing in close proximity to the DNA (see details in *SI Methods*). In order to define what constitutes the “outside” region, we established a criteria based on the DNA (number) density. Specifically, we considered a region to be outside the nucleoid if its DNA density was below 0.1 (on a scale of 0 to 1, with 1 representing the maximum density). Consequently, if a region with a high (number) density of bound ribosomes had a DNA density below 0.1, the ribosomes in that region were classified as excluded. Therefore, regions with low DNA density were considered “outside” the nucleoid, while those with high DNA density were regarded as “inside” (as illustrated in Figure 7a and 7b). Using this definition, we calculated the percentages of free and bound ribosomes present inside and outside the nucleoid.

**FIG. 7.**
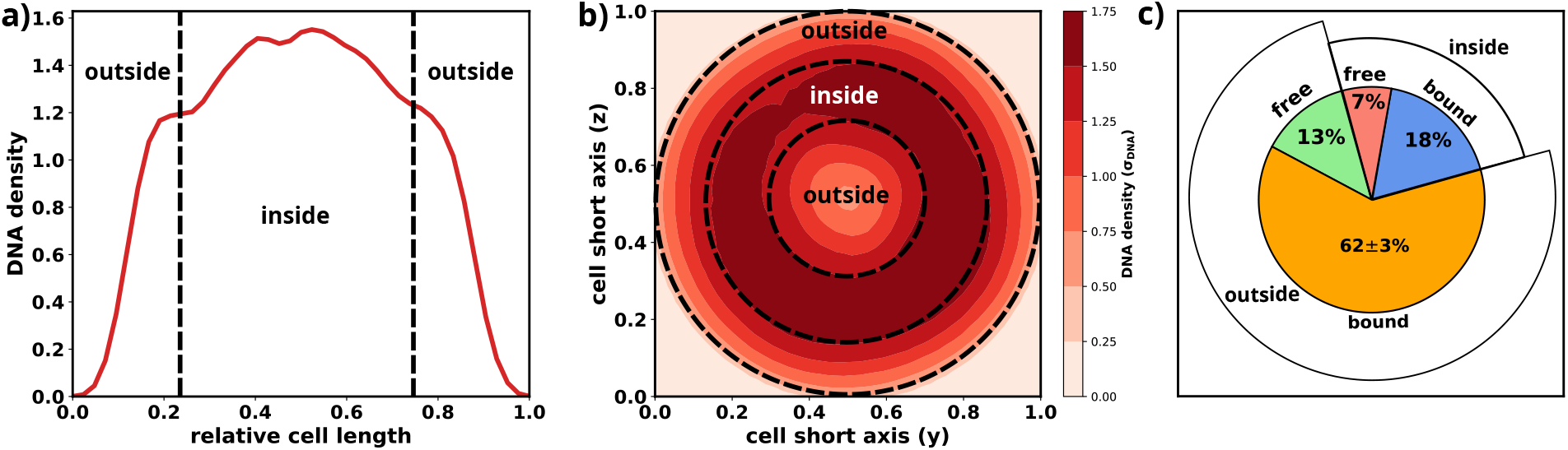
**a)** The chromosome linear density demarcating the regions which can be considered as outside or inside the nucleoid. **b)** A 2D heatmap of the chromosome density demarcating the regions which can be considered as inside or outside the nucleoid. **c)** A pie chart showing the percentage of free and bound ribosomes present inside or outside the nucleoid.

From Figure 7c, it is evident that a relatively small proportion of bound ribosomes (*∼*18%) is found in regions characterized by high DNA density, indicating their presence “inside” the nucleoid. This suggests that the majority of bound ribosomes are indeed excluded from the nucleoid. However, we must note that even bound ribosomes occupying the central void are technically excluded from the nucleoid, although this may not be immediately apparent without a three-dimensional analysis. Therefore, we argue that our model demonstrates a differential tendency in the exclusion of free and bound ribosomal subunits, which can be attributed to their respective sizes.

Free subunits, with dimensions of 14 nm for 30S and 17 nm for 50S, are considerably smaller than the mesh size of the chromosome (50 nm). As a result, they can readily diffuse inside the nucleoid. On the other hand, 70S ribosomal subunits exist as a polymer, with dimensions of approximately 54 1 nm, comparable to the size of the chromosome mesh. Consequently, bound ribosomes face greater difficulty in diffusing into the nucleoid compared to free subunits. However, as demonstrated in a previous section, bound ribosomes can exhibit weak diffusion into the nucleoid through a worm-like movement, leading to a limited population of them being present inside the nucleoid, as depicted in Figure 7c.

Finally, we contemplate the significance behind the exclusion of bound ribosomes from the nucleoid. A 1D reaction-diffusion model, incorporating a wealth of experimental data, has previously predicted the localization of synthesized mRNA predominantly at the endcaps[52]. In the 1D context, the end-caps are considered to be outside the nucleoid, and by extension, their 3D interpretation implies that synthesized mRNA is excluded from the nucleoid. This observation can be reasonably explained by the exclusion of bound ribosomes from the nucleoid. Bound ribosomes represent 70S subunits associated with mRNA during translation. While the formation of bound ribosomes can occur within the nucleoid, nutrients are abundant near the cell wall. Consequently, a substantial portion of “70S-bound mRNA” or polysomes is excluded from the nucleoid, located toward the cell periphery where the availability of nutrients is high, facilitating optimum protein synthesis.

Regarding the small number of ribosomes present in the void, we propose that their role is to translate mRNA molecules that have been excluded from the nucleoid into the void, thus maximizing the utilization of synthesized mRNA. In summary, our data-integrated model not only captures multiple aspects of chromosome topology and organization but represents a pioneering approach that optimizes chromosome-ribosome interactions based on experimental linear densities. This model provides novel insights into the localization of ribosomes within the nucleoid, where a minority of them can also reside within an internal void.

## METHODS

In an attempt to resolve the spatial organization of the ribosomes in *E. coli.*, we build a basic computer model of the bacterial cytoplasm corresponding to medium growth condition (WT *E. coli* cells grown in M9 minimal media at 30 C). The model cytoplasm consisted of a circular chromosome and large number of bound and free ribosomes. The details of the data-integrated chromosome model has been reported by us in a recent article. Here we first provide a brief description of the chromosome model, the details of which has been recently published in a recent article[18]. Then we describe our modeling approach for ribosomes.

### Integrated model of *E.coli* chromosome

We have used multiple experimental data to model the chromosome as a hyper-branched polymer with a circular backbone. We first started with a completely circular chromosome topology. We next tried labeling the chromosome regions into Plectoneme Abundant and Plectoneme Free Regions (PARs and PFRs). We determined the labels by using RNA-Seq signals[12] and that in a genome equivalent of DNA there should be 43*±*10 PARs as described in details in a previous study[18]. Using the labels we generate a hyper-branched polymer with a circular backbone. We then confine the initial conformation in a capsule whose cell dimensions are equivalent to experimentally observed cell dimensions: 3.05 µm in length and 0.82 µm in diameter. We then add ribosomes, both bound and free, in 80:20 ratio[14] into the confinement. This serve as initial configuration for molecular dynamics. We next add Hi-C contact probabilities into the chromosome architecture as distance restraints maintained using harmonic springs as described in the previous study[18]. We then perform molecular dynamics simulations, as described in a later section, to obtain an ensemble to conformations.

### Modeling the cytoplasm of *E. coli*

The *E. coli* cytoplasm is composed of a multitude of biomolecules out of which ribosomes stand out as one of the most abundant and important moieties. Not only are they required for translation of mRNA but they also act as macromolecular crowders for the chromosome thereby helping the bacteria in efficiently packing its DNA inside the small volume its cell has. We model the ribosomes as spherical particles of masses and sizes commensurate with experimental measurements[24, 47](Table III).

**Table III.** Parameters for ribsosomes

| particle type | mass ($m_0 = 0.324 \text{ MDa}$) | size ($\sigma_0 = 68.14 \text{ nm}$) |
| --- | --- | --- |
| DNA | 1 | 0.464 |
| 30S ribosome | 2.62 | 0.205 |
| 50S ribosome | 4.19 | 0.249 |
| 70S ribosome | 7.12 | 0.307 |

The number of ribosomes to be incorporated, per experimental reports[27], were 21000 out of which 80% were 70S subunits and the rest were 30S and 50S subunits[14]. In our current work, we maintained an equal amount of 30S and 50S subunits while all the 70S subunits formed polysomes where each polysome was a polymer of 13 copies of 70S subunits. The exact numbers are 2102 30S subunits, 2102 50S subunits and 1292 13-mer polysomes. The intra-ribosome interactions have been modeled as purely repulsive and DNA-ribosome interactions have been optimized. We have discussed these interactions in next section.

### Interaction potential of the system

The complete interaction potential of the system is summarized as below:

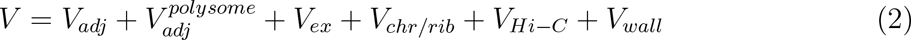

we further elaborate on each term in the upcoming text.

The adjacent beads of the polymer model are bound via harmonic springs defined by Eq-3.

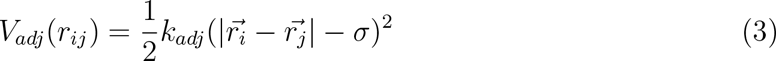

where V*_adj_*(r*_ij_*) is the potential that tethers the adjacent beads of the polymer and k*_adj_* is the stiffness of springs connecting the adjacent beads of the polymer. 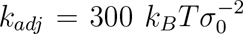 has been used in this study.

The adjacent beads of a polysome are bound by similar type of springs, however much stiffer[14]. It is given by Eq-4.

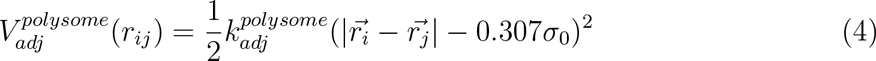

where 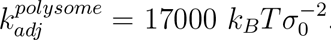.

DNA-DNA and ribosome-ribosome interactions have been modeled as purely repulsive in nature. They have been defined via Eq-5. These interactions are short ranged and are mainly excluded volume interactions.

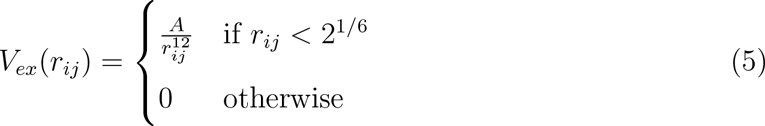

The values of A for different particle pairs (DNA-DNA, polysome-polysome, polysome-ribosome and ribosome-ribosome) have been reported in Table-S1.

However, DNA and ribosome interactions are also not volume exclusion interactions but comprises of the complete Lennard-Jones interaction potential, on the contrary to a previous model[14]. We have defined the potential in Eq-6.

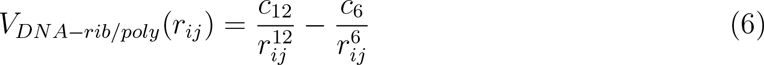

c_6_ and c_12_ have been optimized to reproduce experimental linear density profiles of DNA and ribosomes which have been reported in Table-S1.

The Hi-C contact probabilities have been integrated into the model by first converting them into distances using 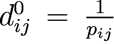. To enforce these distances in the model, we introduce harmonic springs whose spring constants are determined using the equation 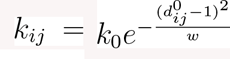. w is a parameter which needs to be optimized and has been optimized in a previous study[18] and 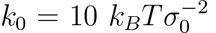 has been used for this study. However, the resolution of the current model is much higher than that of the Hi-C data (10 times higher for the current model). Therefore we incorporate the distance restraints on every fifth bead at an interval of 10. These beads have been labeled as “Hi-C beads”. These beads are bound via harmonic springs which is given by Eq-7.

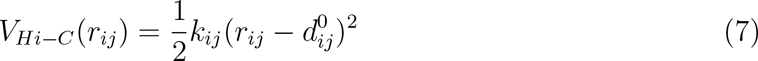

Finally we have also incorporated a confinement potential that acts inwards whenever a particle tries to cross the it to mimic the cell wall. It potential is given by Eq-8.

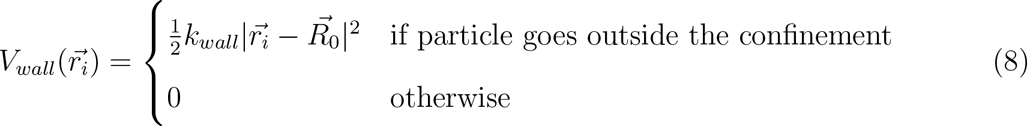

where 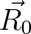 is the center of confinement and 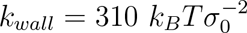 has been used.

### Simulation protocol

We generate 32 random, independent initial configurations inside the confinement satisfying the structural experimental data[12, 42] which we successfully integrate in a polymer model via its architecture. The distances among different regions of the chromosomes are not as per the Hi-C contact probability matrix in the initial conformations of the chromosome. A required number of ribosomes and polysomes[46] are then inserted to obtain a mixture of chromosome and ribosomes which serve as initial configurations.

Using the generated initial configuration, first an energy minimization is run with a force tolerance criteria of 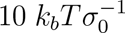. Next molecular dynamics simulations is performed using the Velocity-Verlet algorithm for 3*×*10^6^ steps using a time-step of 0.002 τ*_MD_* at k*_B_*T = 1 where 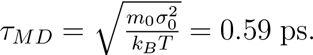. The simulations have been performed via GROMACS 5.0.6[53, 54] which we have modified to include a spherocylindrical confinement. For analysis we have used the last 1000 frames of the generated trajectories.

## Supporting information

Supplemental method, results, figures and table

## ACKNOWLEDGMENT

This work was supported by computing resources obtained from shared facility of TIFR Centre for Interdisciplinary Sciences, India. We acknowledge support of the Department of Atomic Energy, Government of India, under Project Identification No. RTI 4007. We also thank Dr. Sonisilpa Mohapatra for sharing the linear density data published in[27].

## Notes

### Competing Interest Statement

The authors have declared no competing interest.

### Summary of Updates

The abstract has been slightly updated.

