## Supplemental method, results, figures and table for "On the Spatial Positioning of Ribosomes around chromosome in *E. coli* Cytoplasm"

### Supplementary Information

#### METHODS

##### Calculation of size of beads and the simulation length scale

Let the packing fraction of DNA inside the cell be $f$. Let the length of the bacterial cell including the end caps be $L$ and the diameter be $d$. Let $l = L - d$. Therefore volume of the cell is given by

$$V_{cell} = \frac{\pi d^2}{2}(\frac{d}{3} + \frac{l}{2}) \tag{1}$$

Let the size of each bead be $\sigma$. Let there be $N$ beads where $N = \frac{4.64 \times 10^6}{res}$ where $res$ is the resolution of the model in base pairs. The volume of each bead is given by

$$V_{bead} = \frac{\pi \sigma^3}{6} \tag{2}$$

Therefore, $f \times V_{cell} = N \times V_{bead}$. Thus the size of each bead ($\sigma$) is given by

$$\sigma = d\sqrt[3]{\frac{1}{N}(1 + \frac{3l}{2d})} \tag{3}$$

Using $f = 0.1$, $res = 500$ bp, $L = 3.05$ $\mu m$, $d = 0.84$ $\mu m$, $\sigma = 31.61$ nm. However to keep simulation boxes similar to our previous studies[1, 2], we have used the simulation length scales ($\sigma_0$) as 68.1 nm. Therefore $\sigma = 0.464$ $\sigma_0$, $L = 44.71$ $\sigma_0$ and $d = 12.32$ $\sigma_0$.

##### Force-field for the chromosome

The mass of each DNA bead has been set to $1m_0$ which is equal to 0.324 MDa[3] Therefore the complete force field for the chromosome is given by

$$V(r) = V_{adj}(r_{ij}) + V_{ex}(r_{ij}) + V_{wall}(\vec{r_i}) + V_{Hi-C}(r_{ij}) \tag{4}$$

$$V_{adj}(r_{ij}) = \frac{1}{2}k_{adj}(|\vec{r_i} - \vec{r_j}| - \sigma)^2 \tag{5}$$

where $V_{adj}(r_{ij})$ is the potential that binds adjacent beads of the polymer and $k_{adj}$ is the force constant of the springs connecting adjacent beads of the polymer. $k_{adj} = 300$ $k_B T \sigma_0^{-2}$ has been used.

$$V_{ex}(r_{ij}) = \frac{A}{r_{ij}^{12}} \quad (6)$$

where $V_{ex}(r_{ij})$ is the volume exclusion potential that prevents beads of the polymers from overlapping. We have used $A = 4.0\ k_BT\sigma_0^{12}$ for our model.

$$V_{wall}(\vec{r_i}) = \begin{cases} \frac{1}{2}k_{wall}|\vec{r_i} - \vec{R_0}|^2 & \text{if particle goes outside the confinement} \\ 0 & \text{otherwise} \end{cases} \quad (7)$$

To mimic the cell wall of the bacteria, we have used a capsule shaped confining potential $V_{wall}(\vec{r_i})$. It acts only when a particle goes outside the potential and forces the particle back towards the centre of the confinement which is given by $\vec{R_0}$. $k_{wall} = 310\ k_BT\sigma_0^{-2}$ has been used for this study.

$$V_{Hi-C}(r_{ij}) = \frac{1}{2}k_{ij}(r_{ij} - d_{ij}^0)^2 \quad (8)$$

where $V_{Hi-C}(r_{ij})$ is the potential that maintains proper distances among the Hi-C beads obtained from experimental contact probability matrix and $k_{ij} = k_0 \exp[\frac{-(d_{ij}^{(0)} - \sigma_0)^2}{w}]$.
$k_0 = 10\ k_BT\sigma_0^{-2}$ has been used for this study.

$w$ in Eq-8 needs to be optimized for proper reproducibility of the contact probability matrix from simulations. We have obtained an optimum value of $w = 1.0$. The comparison between the simulated and experimental contact probability matrices have been shown in Figure 1.

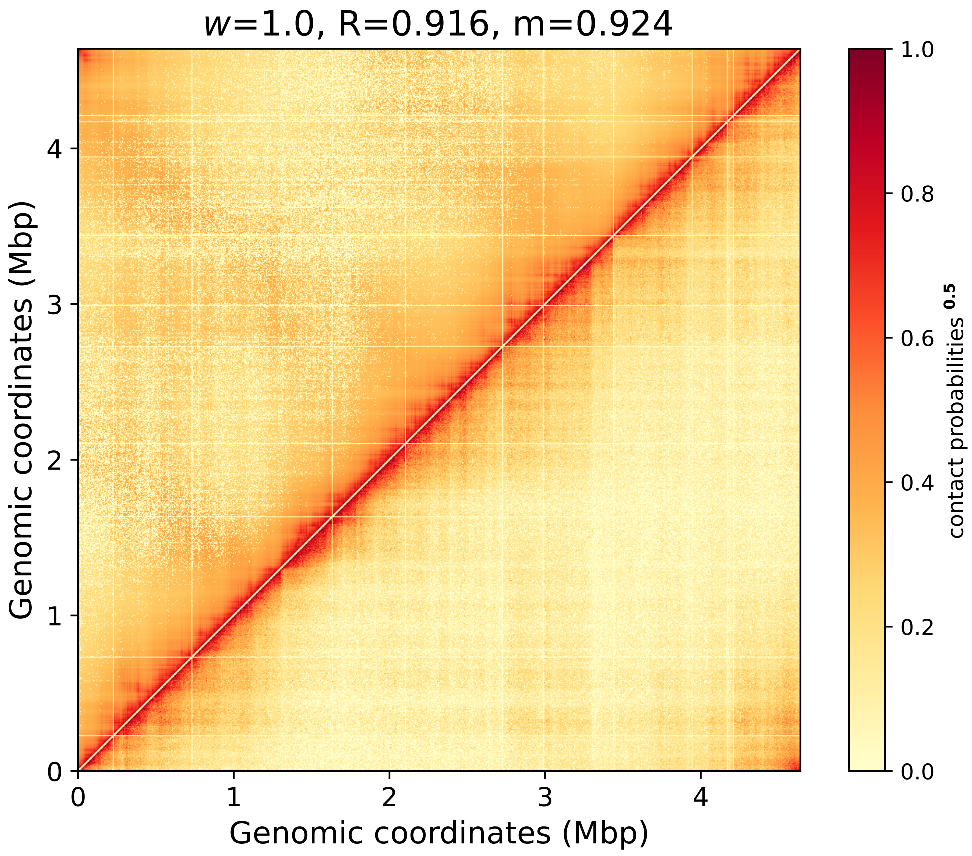


FIG. 1. Comparison of experimental and simulated contact probability matrix for $w = 1.0$. Upper triangle represents probabilities obtained via simulations while the lower triangle denotes the experimental matrix. The values have been enhanced (by using $\sqrt{\text{contact probabilities}}$) for a better visual comparison.

##### Modeling ribosomes and polysomes

Ribosomes comprise of 30S and 50S sub-units in *E. coli*. The 30S and 50S together can form a 70S ribosome complex, which polymerizes to form polysomes. Polysomes are the function form of ribosomes that help in translation of mRNA into proteins. Each 30S, 50S and 70S monomeric subunit have been modeled as spherical particles with masses and sizes (in terms of $\sigma_0$) as provided in Table-**??**. The polysomes have been modeled as a 13-mer of 70S subunits[4]. We have incorporated 21000 particles belonging to 30S, 50S and 70S. Out of them 1292 are 30S, 1292 are 50S and 18416 particles (=2102 polysome chains) are 70S[5].

##### Force-field for the ribosomes and polysomes

The ribosomes and polysomes have interactions with themselves, with the DNA and with the wall ($V_{wall}(\vec{r_i})$). The $V_{wall}$ confines ribosomes and polysomes in the cell wall and has the same form as Eq 7.

The ribosomes and polysomes with themselves have purely repulsive interactions which has the same form as Eq 6. The value of $A$ between ribosomes have been provided in Table-S1.

The monomeric beads of the polysomes are tightly bound together via a harmonic potential which has the same form as Eq 5 and given by

$$V_{adj}^{polysome}(r_{ij}) = \frac{1}{2}k_{adj}^{polysome}(|\vec{r_i} - \vec{r_j}| - 0.307\sigma_0)^2 \tag{9}$$

where $k_{adj}^{polysome} = 17000\ k_B T \sigma_0^{-2}$.

##### Non-bonded interactions among particles

DNA-DNA, ribosome-ribosome, polysome-polysome and ribosome-polysome interactions are purely repulsive and have the same form as of Eq 6. Values of $A$ for respective pairs have been provided in Table-S1.

However, complete Lennard-Jones potential has been considered for the ribosomal or polysomal interaction with DNA and is given by

$$V_{DNA-rib/poly}(r_{ij}) = \frac{c_{12}}{r_{ij}^{12}} - \frac{c_6}{r_{ij}^6} \tag{10}$$

where $c_6$ and $c_{12}$ have been tuned to match the simulated and experimental linear density profiles of DNA and ribosomes.

Details of $c_6$ and $c_{12}$ have been provided in Table-S1.

##### Interaction of probes with the particles in the system

Let the size of the probe be $\sigma_{probe}$, size of a 30S ribosome be $\sigma_{30S}$, size of a 50S ribosome be $\sigma_{50S}$, size of a 70S ribosome be $\sigma_{70S}$ and size of a bead of the polymer representing the chromosome be $\sigma_{DNA}$. Except for $\sigma_{probe}$, all other sizes have been provided in Table-**??**. The interaction of a probe with itself and any other particle is purely repulsive and is given by

$V_{probe}(r) = \frac{A_{probe}}{r^{12}}$ where $A_{probe} = 4 * \sigma'^{12}$ and $\sigma'$ is the effective size of the particles given by $\sigma' = \sqrt{\sigma_{probe}\sigma_{30S/50S/70S/DNA}}$. We have used probe sizes of 25 nm, 40 nm, 45 nm 47.5 nm, 50 nm, 52.5 nm, 55 nm and 60 nm.

Therefore if we wanted to calculate the interaction potential between a probe and a 30S ribosome particle, then $\sigma' = \sqrt{\sigma_{probe} * \sigma_{30S}}$ and $V_{probe/30S}(r) = \frac{4\sigma'^{12}}{r^{12}}$. If the probe size is 50 nm, $\sigma_{probe} = \frac{50}{68.14} = 0.733\sigma_0$ and $\sigma' = 0.387\sigma_0$. The corresponding $A_{probe} = 4 * (0.387\sigma_0)^{12} = 4.6 \times 10^{-5}\sigma_0^{12}$.

**Calculation of 3D densities for ribosomes and chromosome**

The percentage was obtained by first placing the cell on a 3D grid. Each cell is called a voxel. The width of the side of each cubic voxel is 0.7 nm. We then calculated the density of DNA in each voxel. Next we determined the position of each ribosome subunit and to which voxel each belonged to. However we considered ribosome subunits present only in the cylindrical region of the cell, i.e. we did not consider ribosomes that populate the end caps. Ribosomes present in voxels with DNA density lower than 0.01 have been classified as being "outside" the nucleoid, i.e. excluded. The rest have been labeled as "inside" the nucleoid

#### SUPPLEMENTARY TABLES

TABLE S1. All non-bonded interaction parameters.

| particle-1 | particle-2 | $c_6$ $(k_B T \sigma_0^6)$ | $A/c_{12}$ $(k_B T \sigma_0^{12})$** |
|---|---|---|---|
| DNA | DNA | —— | 1.0 |
| 30S ribosomes | 30S ribosomes | —— | $2.236\times10^{-8}$ |
| 50S ribosomes | 50S ribosomes | —— | $2.298\times10^{-7}$ |
| 70S ribosomes | 70S ribosomes | —— | $2.901\times10^{-6}$ |
| 30S ribosomes | 50S ribosomes | —— | $7.167\times10^{-8}$ |
| 50S ribosomes | 70S ribosomes | —— | $8.164\times10^{-7}$ |
| 30S ribosomes | 70S ribosomes | —— | $2.547\times10^{-7}$ |
| DNA | 30S ribosomes | $3.459\times10^{-4}$ | $2.990\times10^{-8}$ |
| DNA | 50S ribosomes | $3.459\times10^{-4}$ | $9.587\times10^{-8}$ |
| DNA | 70S ribosomes | $3.459\times10^{-4}$ | $3.406\times10^{-8}$ |

**For same particle and all ribosome-ribosome interactions, there no $c_6$ and only $A$.

For DNA-ribosome interactions only should consider $c_6$ and $c_{12}$.

#### SUPPLEMENTARY RESULTS

##### The model cytoplasm represents a poor solvent with a 50 nm mesh size

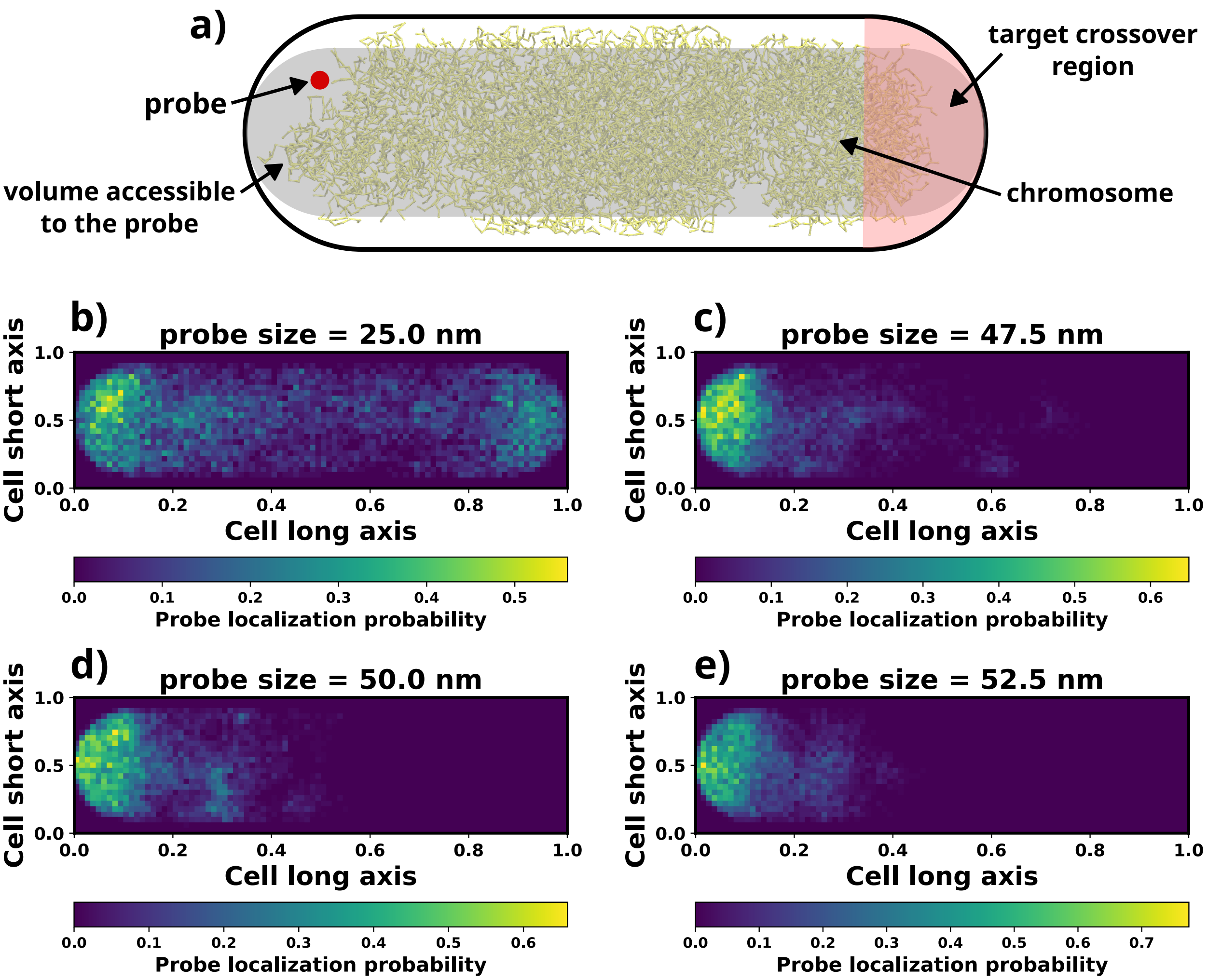


FIG. 2. **a)** A schematic showing how the mesh was size was determined using a probe (red). The volume accessible to the probe has been highlighted in grey. The target region where the probe needed to end up for it to considered as being able to pass through the nucleoid has been shown in light red. Locations a probe of size **b)**25 nm **c)** 47.5 nm **d)** 50 nm **e)** 52.5 nm visited. The colorbar represents the probability with which the probe visited that particular location. We can see that the probe cannot enter the nucleoid starting from $\sim$50 nm.

From our simulations, we observed that the chromosome adopts a completely collapsed conformation inside our confinement. The earlier report and the above observation motivated us to estimate the Flory exponent from our simulations using the Eq-11 proposed by Xiang et al.[6].

$$\xi = \frac{\sqrt{3}}{6} b^{\frac{\nu-1}{3\nu-1}} \left( \frac{18\sqrt{3} M_W}{\pi c N_A I_{bp}} \right)^{\frac{\nu}{3\nu-1}} \tag{11}$$

where $\xi$ is the mesh size of the chromosome which will be determined later in this section. $M_W$ is the average molecular weight of the chromosome which is 650 g/mol[3], $N_A$ is Avogardo's number, $I_{bp}$ is size of a base-pair[7, 8] and $c$ is the concentration of DNA. Therefore for estimation of the Flory exponent, we first need to obtain the values of mesh size ($\xi$) and concentration of DNA ($c$) which we determine next.,

*Determination of mesh size:* To estimate the mesh size of the chromosome[6], we inserted "probes" of different diameters at one end of the *E. coli* cell (Figure 2a). The interactions of the probe with the rest of the particle is given by $V_{probe}(r) = \frac{A_{probe}}{r^{12}}$ where $A_{probe} = 4\sigma'^{12}$, $\sigma' = \sqrt{\sigma_{probe}\sigma_{particle}}$ and $\sigma_{particle}$ is provided in Table-S1. More details can be found in the *SI Methods* section. For estimation of the mesh size, we have used probes of sizes 25 nm, 47.5 nm, 50 nm and 60 nm.

Upon successful insertion of a probe, we confined the probe to a volume such that if it has to cross over to the other end of the cell, it has to pass through the chromosome (Figure 2a). Next we let the system evolve. Once we have evolved the system for a sufficient number of steps ($3 \times 10^6$ steps), we calculated a 2D-histogram of the probe inside the cell (Figure 2b-e). If the probe is able to cross over to the other side of the cell, then we should observe sufficient probability of the probe on both cell poles which can be seen from Figures 2b for the 25 nm probe. However, as we kept increasing the probe size, we saw lesser probability of the probe being present on the side opposite to its starting side. We hypothesized that if the probe size is greater than or equal to the mesh size, it would not be able to cross over at all. We performed four independent runs with each seed and averaged the 2D probability density for all sizes over all independent runs (Figure 2b-e). What we observed is that for probe size = 50 nm, the cross-over to the other side has completely stopped as it cannot penetrate through the mesh that the chromosome has made. Therefore we estimated that the mesh size of the chromosome to be ∼50 nm which is also the experimentally estimated value[6].

*Determination of concentration of DNA:* As the chromosome has a fixed number of base pairs and the average molecular weight of a base pair in a dsDNA is known, the mass of the chromosome ($M_{chr}$) is fixed and can be calculated as $M_{chr} = \frac{650 \times 4.64 \times 10^6}{N_A}$ g where $N_A$ is the Avogadro's number. The concentration ($c$) can then be calculated as $c = \frac{M_{chr}}{V_{chr}}$ where $V_{chr}$ is the volume occupied by the chromosome. As the chromosome shape inside the cell is an ellipsoid, we determined its volume using Eq-12.

$$V_{chr} = 4\pi\sqrt{3\lambda_1\lambda_2\lambda_3} \tag{12}$$

where $\lambda_1$, $\lambda_2$ and $\lambda_3$ are the eigenvalues from the gyration tensor of the chromosome. Using the above equation we estimate that that volume occupied by the chromosome is 0.53 $\mu m^3$. Therefore the concentration we obtain is 9.5 mg/mL which is close to the experimentally estimated concentration of 7 mg/mL.

The Kuhn length ($b$) of a polymer is given by twice its persistence length ($l_p$). For our chromosome model, since we have not incorporated any angular potentials, the persistence length is equal to the distance between two adjacent beads which is 31.66 nm. Thus, for our model, $b = 63.32$ nm which is close to the experimentally determined value of 60 nm[6].

Upon incorporating $c = 9.5$ mg/mL, $b = 63.32$ nm and $\xi = 50$ nm into Eq-11 and solving $\frac{\sqrt{3}}{6} b^{\frac{\nu-1}{3\nu-1}} \left(\frac{18\sqrt{3}M_W}{\pi c N_A I_{bp}}\right)^{\frac{\nu}{3\nu-1}} - \xi = 0$ iteratively for $\nu$ provides us with $\nu = 0.31$ having the value closest to zero (Figure S3) with an error of 1.07 nm (2.15 % error).

Thus we observe that the Flory exponent we obtained from our simulations matches closely with that of the experimentally obtained value. It also tells us that for our model, the interactions among different particles and among different regions of the chromosome capture the effect of the cytoplasm being a bad solvent for the chromosome very efficiently.

**Control simulations to prove that the void inside the nucleoid is not an artifact**

To prove that the void inside the nucleoid is not an artifact of the simulations, we perform a set of simulations where there are no ribosomes present. The two-dimensional heat-map of the DNA density in such a scenario (Figure 3) still shows low DNA density at the centre thereby confirming that the central void exists independent of the presence of ribosomes. We think that this has to do with the fact that the chromosome conformation is mainly maintained by the multiple constraints we have incorporated into our model by integrating it with RNA-seq and Hi-C data. The presence of ribosomes only act as weak perturbations to the rather stringent and highly constrained chromosome, thus rendering them a bit more flexible and being able to vary its conformation slightly more than what it could have without the ribosomes. Moreover the central void is agnostic of nature of chromosome-ribosome interaction: control simulation in which the ribosomes are made *inert*, also retains the void in the centre of the cytoplasm.

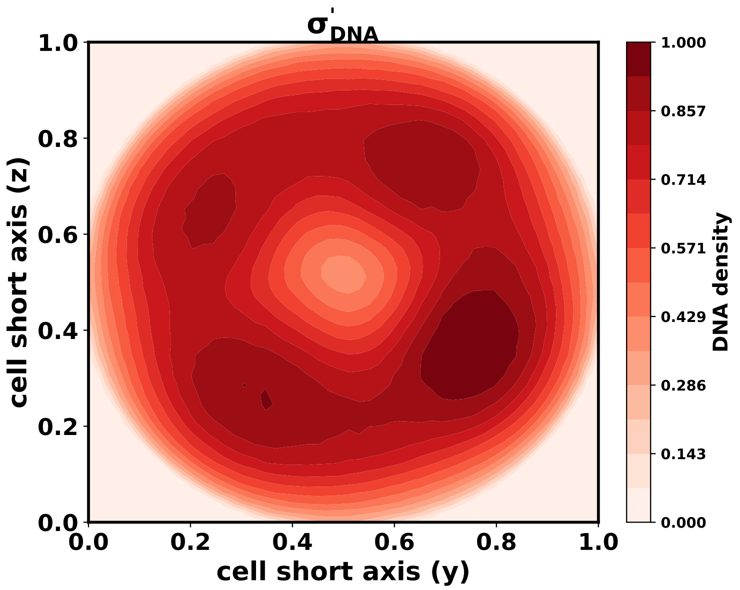


FIG. 3. **a)** Contour plot showing the DNA density ($\sigma^{'}_{DNA}$) along the radial directions for a simulation set without any ribosome.

*The void is not an artifact of initial configuration generation*: However one can still argue that the ribosome density inside the void is due trapped ribosomes caused by the initial configuration since they were prepared in the following way: i) the chromosome architecture is first generated. ii) it is put inside the confinement by turning on all the potentials except the Hi-C interactions and performing an energy minimization. iii) ribosomes are added to the system. iv) Hi-C interactions are now turned on and molecular dynamics simulations are performed.

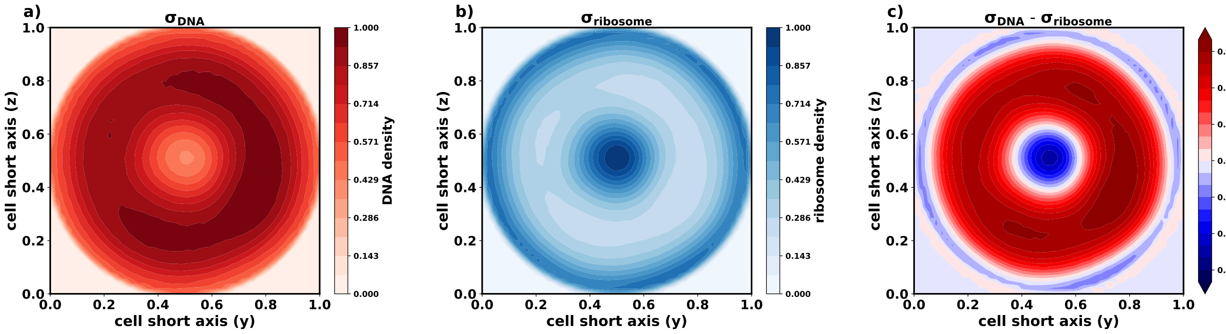


FIG. 4. For late insertion of ribosomes into the simulations, **a)** Contour plot showing the DNA density ($\sigma_{DNA}$) along the radial directions. **b)** Contour plot showing the ribosome density ($\sigma_{ribosome}$) along the radial directions. **c)** Contour plot showing the difference in DNA and ribosome densities ($\sigma_{DNA}$ - $\sigma_{ribosome}$) along the radial directions.

Therefore to mitigate the issue and have more confidence in the current results, we ran a new set of simulations where between step-ii and step-iii, an extra step was performed where the Hi-C interactions are turned on and the confined chromosome conformations underwent molecular dynamics simulation for $3 \times 10^6$ steps. The ribosomes and polysome were then added as per step-iii and molecular dynamics simulations were performed for $3 \times 10^6$ more steps as in step-iv. We recalculated the DNA and ribosome 2D densities for these new set of simulations (Figure 4). These densities still show the central void with high ribosome density at the centre. Therefore the ribosome density at the centre is not an artifact of initial configuration generation and the control simulations aid our observation that of ribosomes are present inside the nucleoid's void in high numbers and that the void is an intrinsic feature of the chromosome organization itself.

*The population of the void is chromosome-ribosome interaction agnostic* We next try to investigate whether the interaction we have considered for the chromosome and the ribosomes has helped the ribosomes populate the insides of the nucleoid. For that we considered a fully repulsive potential between the DNA and the ribosomes are given by Eq-13.

$$V_{DNA-rib/poly}(r_{ij}) = \frac{c_{12}}{r_{ij}^{12}} \tag{13}$$

Eq-13 is simply the repulsive part of the full Lennard-Jones potential we have considered during modelling the DNA-ribosome interactions as per Eq-10. We perform a set of molecular dynamics simulation using Eq-13 instead of Eq-10 and recalculate the 2D densities for the chromosome and the ribosomes (Figure S1). We observe that the densities still show a ribosome populated central void which confirms our hypothesis that irrespective of whether the chromosome-ribosome are purely repulsive or not.

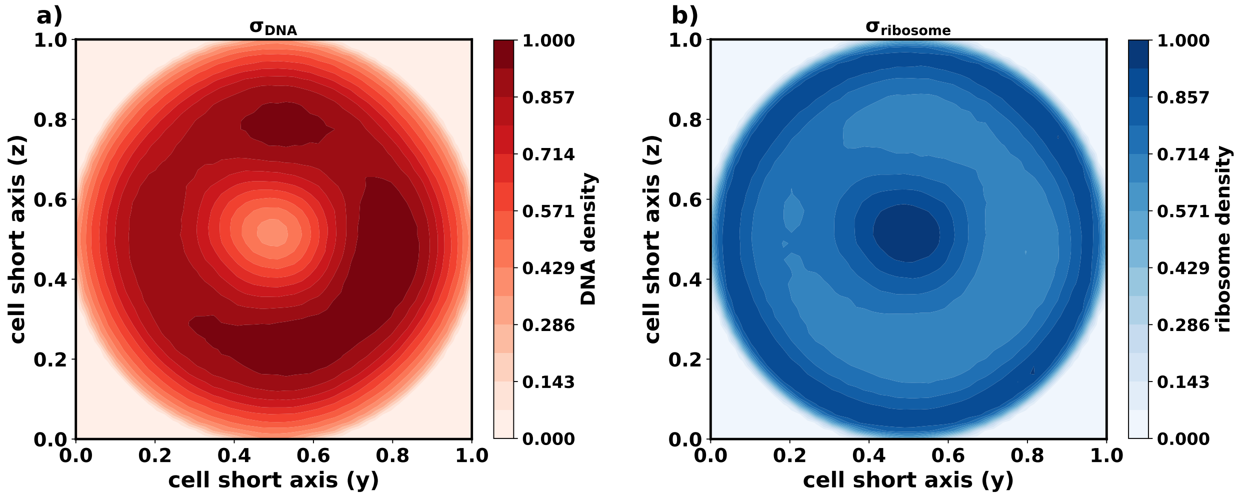


FIG. 5. **a)** 2D DNA densities for purely repulsive chromosome-ribosome interactions. **b)** 2D densities of ribosomes for purely repulsive chromosome-ribosome interactions

**Control simulations proving that polysomes can percolate through the chromosome mesh**

To verify whether polysomes outside the nucleoid can percolate through the nucleoid and access the void at the centre of the cell, we ran a new set of simulations. These simulations started with scenario where all polysomes were initially placed outside the cell confinement (Figure S6). An energy minimization caused them to be placed inside the cell. We then ran molecular dynamics simulations for $3 \times 10^6$ steps and obtained an ensemble of simulation trajectories. Using the last 1000 frames from each trajectory of the ensemble, we recalculate the linear (Figure S7) and the 2D densities (Figure 6) for the DNA and the polysomes. We observe that the linear densities from the control simulations match almost exactly with the linear densities shown in Figure 2c and e (also shown in dotted lines in Figure S7).

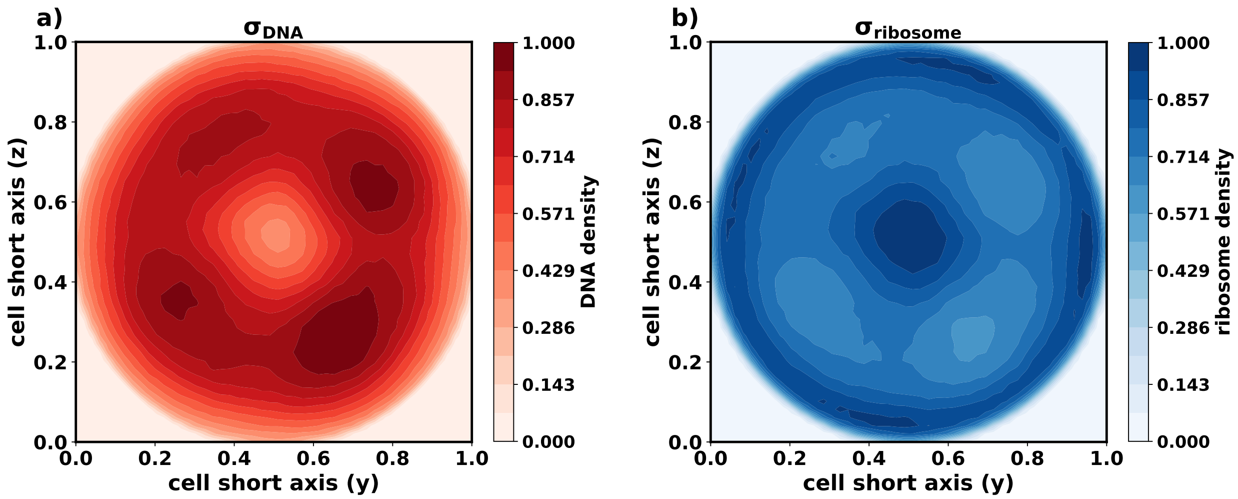


FIG. 6. The following 2D densities are for simulations whose starting conformations were like Figure S6. **a)** 2D density for the chromosome. **b)** 2D densities for the ribosomes.

##### Determining the polysome conformation at different regions of the chromosome

To determine how polysomes outside the nucleoid could gain access to the void inside it, we first determined the effective size of polysomes at different part of the cell cross-section. To that end, we first binned the radial axes into a grid and categorized each polysome into a grid point based on its centre of mass (see *SI Methods* for more details). Once we have categorized them, we calculated the average radius of gyration and average end-to-end distance of the polysomes belonging to a particular grid. We then repeated the calculation for all other grids to obtain a matrix that contained the mean radius of gyration (Figure 7b) and end to end distance (Figure 7c) values along with their standard deviations. We can see that for all bins $R_g$ and $R_{ee}$ do not change much. Thus we can safely assume that almost all polysomes have conformations that are similar in terms of their sizes. Since the polysomes are present in extended conformations, the effective cylindrical volume for them would provide an estimate if they can percolate through the mesh of size 50 nm. To generate the effective cylindrical representation, we calculated the length ($L$), width ($d_x$) and breadth ($d_y$) of each polysome (as shown in Figure S9a). The diameter of the cylinder ($d$) is an average of $d_x$ and $d_y$. We obtain $\bar{L}$ = 99.2 nm and $\bar{d}$ = 59.8 nm (Figure 7d). These values are very close to the mesh size and they, in practice, underestimate the actual percolation propensity by assuming the **flexible** polysome as a **rigid** cylinder. Thus the polysome can easily wiggle through the mesh and is able to, thus, occupy the central void that occurs due to the chromosome's own arrangement.

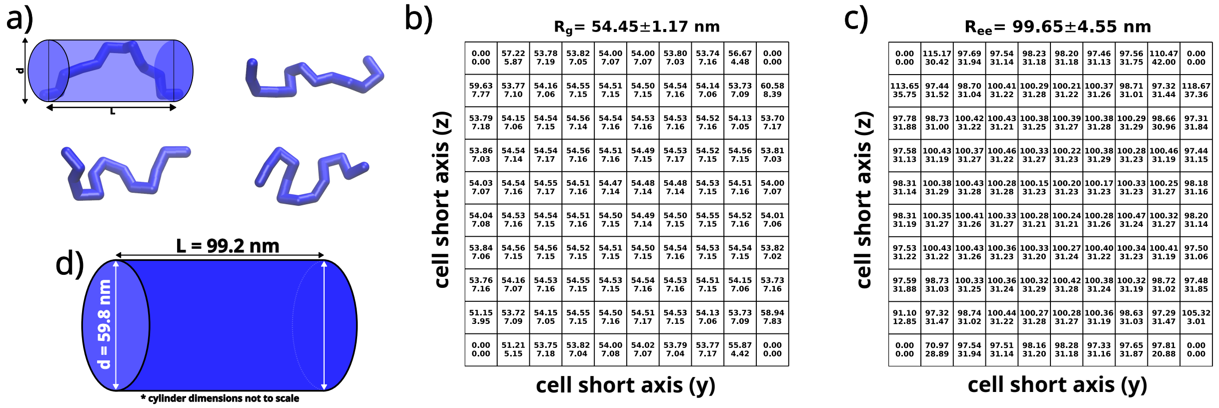


FIG. 7. **a)** Different representative polysome conformations. **b)** each cell dimensions has been divided into 10 bins. Thus the two short axes form a grid/matrix. We determined the mean and standard deviations of **b)** radius of gyrations **c)** end to end distances of polysomes present in each bin which we have reported in the form of the matrices. **d)** Representing a polysome as a cylinder.

To generate the effective cylindrical representation, we calculated the length ($L$), width ($d_x$) and breadth ($d_y$) of each polysome (as shown in Figure S9a). The diameter of the cylinder ($d$) is an average of $d_x$ and $d_y$. We obtain $\bar{L} = 99.2$ nm and $\bar{d} = 59.8$ nm (Figure 7d). These values are very close to the mesh size and they, in practice, underestimate the actual percolation propensity by assuming the **flexible** polysome as a **rigid** cylinder. Thus the polysome can easily wiggle through the mesh and is able to, thus, occupy the central void that occurs due to the chromosome's own arrangement.

#### SUPPLEMENTARY FIGURES

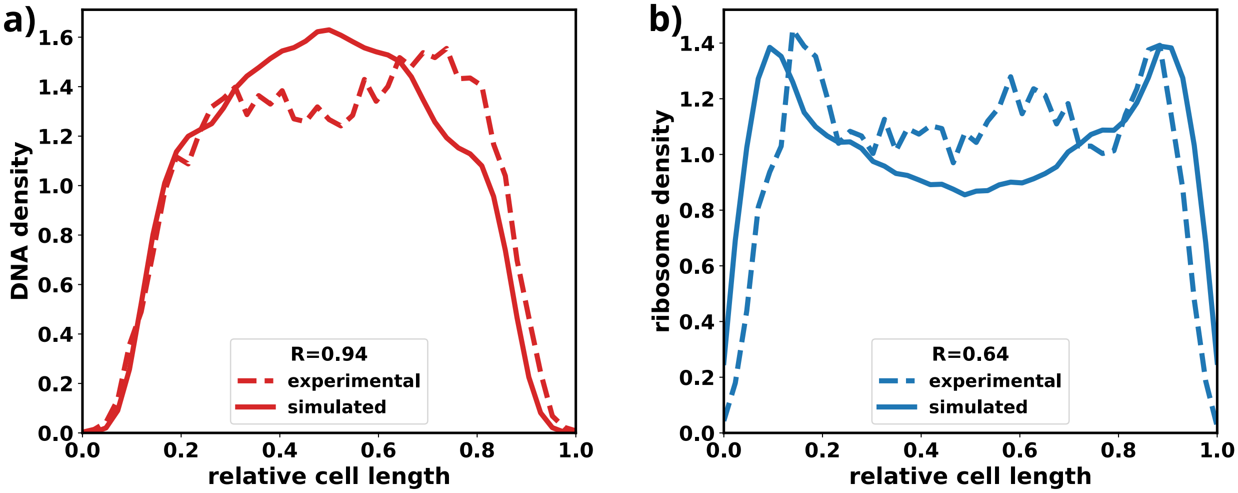


FIG. S1. **a)** Comparison of experimental linear densities for the chromosome (dashed) and linear densities obtained from simulations (bold) where the ribosomes have been modeled as inert crowders.**b)** Comparison of experimental linear densities for ribosomes (dashed) and linear densities obtained from simulations (bold) where the ribosomes have been modeled as inert crowders.

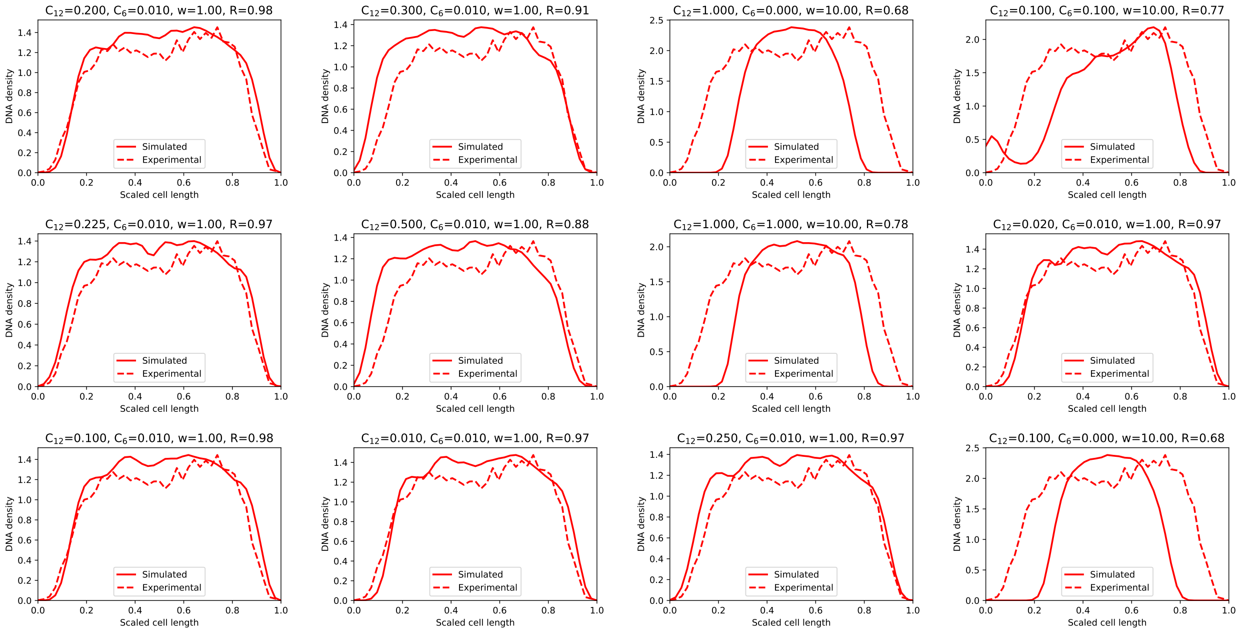


FIG. S2. The DNA densities for different interaction parameter sets.

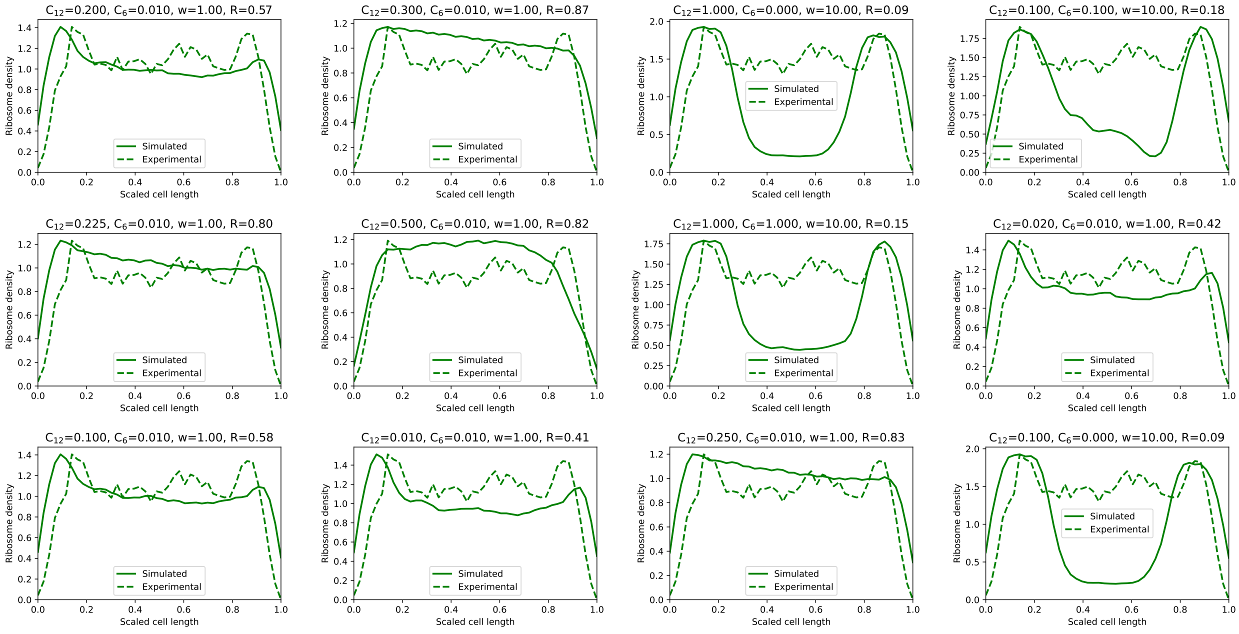


FIG. S3. The ribosome densities for different interaction parameter sets.

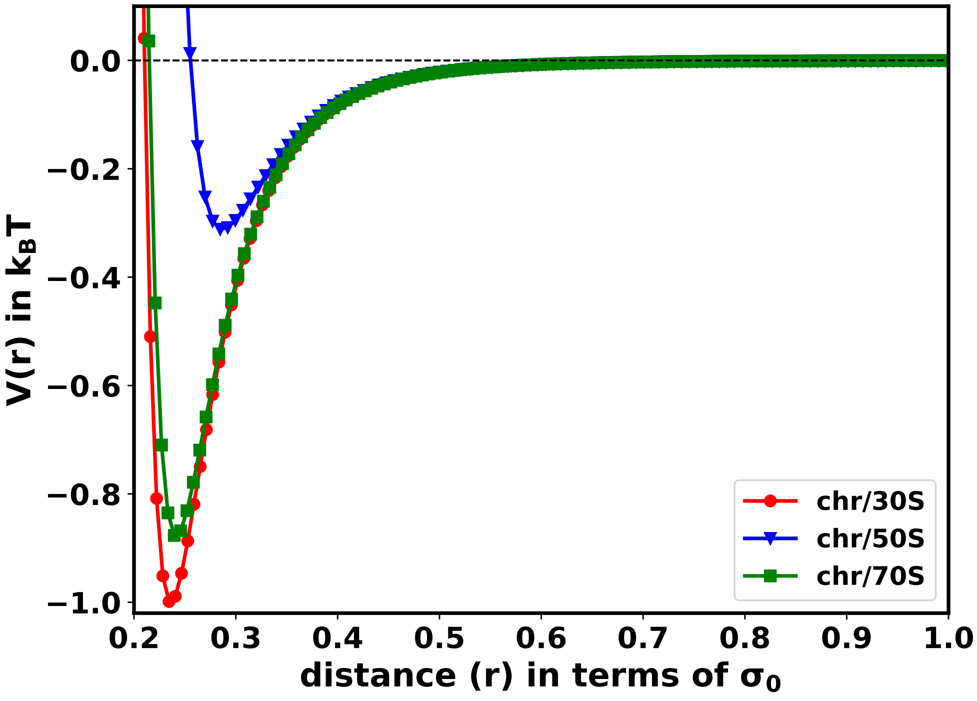


FIG. S4. The chromosome-ribosome interaction potentials for each ribosomal subunit.

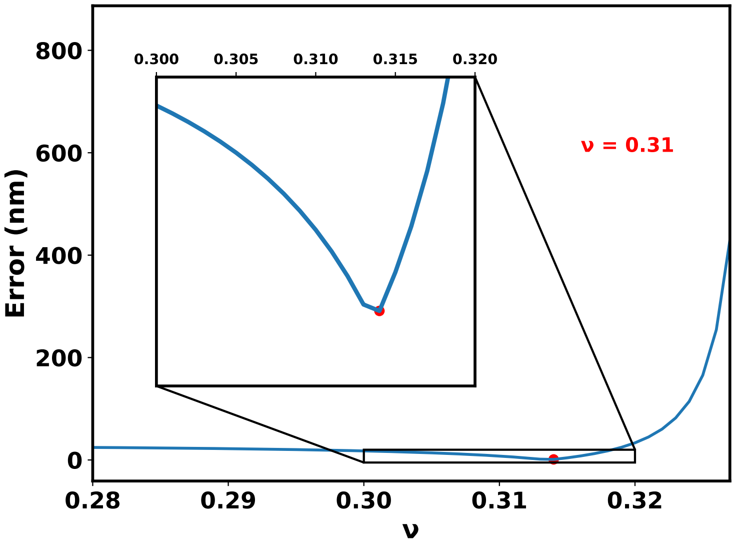


FIG. S5. Error vs. the scaling exponent ($\nu$) in Eq 13.

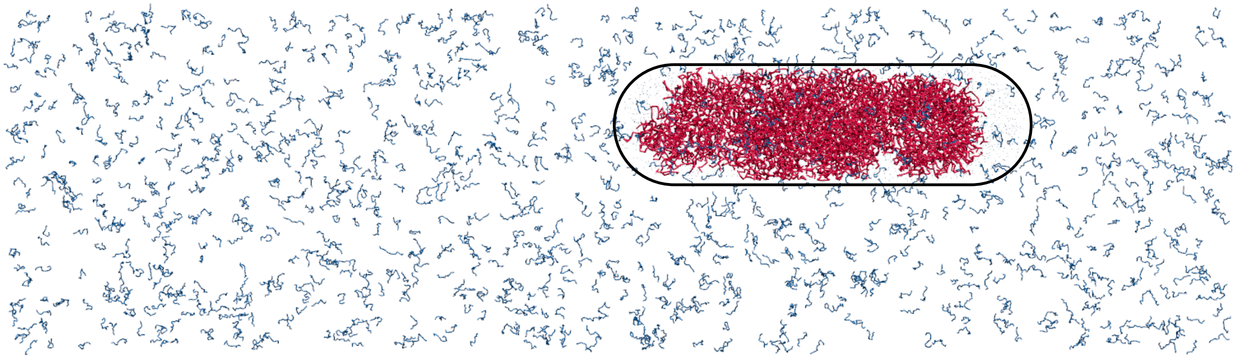

FIG. S6. The initial configuration where all polysomes (blue) were placed outside the cell. The chromosome is represented in red. The black capsule represents the cell wall.

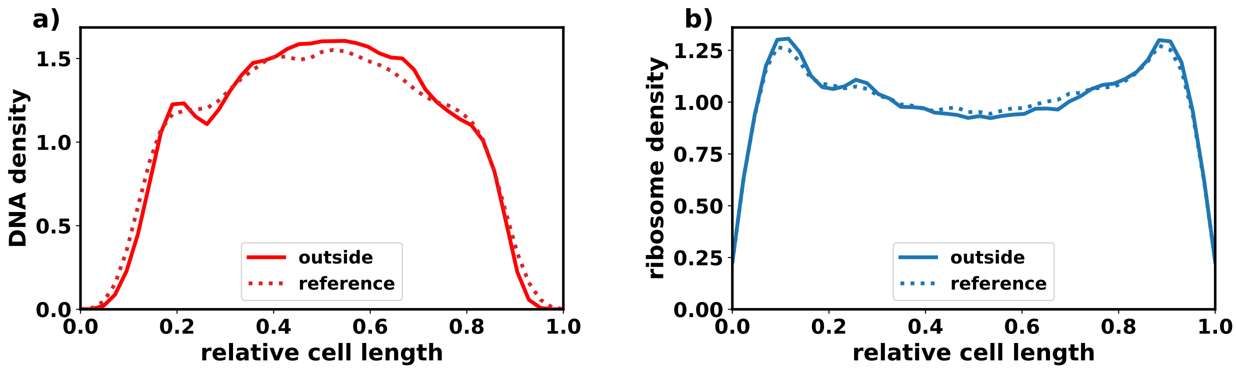


FIG. S7. The bold line represents the linear densities obtained from simulations with initial configuration as shown in Figure S6. The dotted line represents the linear densities shown in Figure 2c and e. **a)** linear densities for DNA **b)** the linear densities for ribosomes.

---

[1] Abdul Wasim, Ankit Gupta, and Jagannath Mondal. A hi–c data-integrated model elucidates e. coli chromosome's multiscale organization at various replication stages. *Nucleic acids research*, 49(6):3077–3091, 2021. 2

[2] Palash Bera, Abdul Wasim, and Jagannath Mondal. Hi-c embedded polymer model of escherichia coli reveals the origin of heterogeneous subdiffusion in chromosomal loci. *Physical Review E*, 105(6):064402, 2022. 2

[3] https://bionumbers.hms.harvard.edu/files/Nucleic%20Acids_Sizes_and_Molecular_Weights_2pgs.pdf. 2, 9

[4] Jagannath Mondal, Benjamin P Bratton, Yijie Li, Arun Yethiraj, and James C Weisshaar. Entropy-based mechanism of ribosome-nucleoid segregation in e. coli cells. *Biophysical journal*, 100(11):2605–2613, 2011. 4

[5] Sonisilpa Mohapatra and James C Weisshaar. Functional mapping of the e. coli translational machinery using single-molecule tracking. *Molecular microbiology*, 110(2):262–282, 2018. 4

[6] Yingjie Xiang, Ivan V Surovtsev, Yunjie Chang, Sander K Govers, Bradley R Parry, Jun Liu, and Christine Jacobs-Wagner. Interconnecting solvent quality, transcription, and chromosome folding in escherichia coli. *Cell*, 184(14):3626–3642, 2021. 9, 10

[7] S Diekmann, W Hillen, B Morgeneyer, RD Wells, and D Pörschke. Orientation relaxation of dna restriction fragments and the internal mobility of the double helix. *Biophysical Chemistry*,

15(4):263–270, 1982. 9

[8] Kensuke Yonemura and Hiroshi Maeda. A new assay method for dnase by fluorescence polarization and fluorescence intensity using dna-ethidium bromide complex as a sensitive substrate. *The Journal of Biochemistry*, 92(4):1297–1303, 1982. 9
